# Rampant transposition following RNAi loss causes hypermutation and antifungal drug resistance in clinical isolates of a human fungal pathogen

**DOI:** 10.1101/2021.08.11.455996

**Authors:** Shelby J. Priest, Vikas Yadav, Cullen Roth, Tim A. Dahlmann, Ulrich Kück, Paul M. Magwene, Joseph Heitman

## Abstract

Microorganisms survive and compete by stochastically acquiring mutations that enhance fitness. Although increased mutation rates are often deleterious in multicellular organisms, hypermutation can be beneficial for microbes experiencing strong selective pressures. Infections caused by *Cryptococcus neoformans* are responsible for ∼15% of AIDS-related deaths and associated with high mortality rates, attributable to a dearth of antifungal drugs and drug resistance. We identified two hypermutator *C. neoformans* clinical isolates in which Cnl1 transposon insertions were responsible for drug resistance. Whole-genome sequencing revealed both hypermutator genomes harbor a nonsense mutation in the RNAi component *ZNF3* and hundreds of Cnl1 elements organized into massive subtelomeric arrays on every chromosome. QTL mapping identified a significant locus associated with hypermutation that included *znf3*. CRISPR-mediated editing of the *znf3* nonsense mutation abolished hypermutation and restored siRNA production. In sum, hypermutation and drug resistance in these isolates results from RNAi loss and a significant burden of Cnl1 elements.

## Introduction

Stochastic mutations and genomic rearrangements provide variation in populations for natural selection to act upon and enable evolution. However, genetic changes are a double-edged sword: too little variation can lead to evolutionary stagnation, while too much can lead to a lethal accumulation of deleterious mutations. Hypermutation, one extreme of this mutational spectrum, can lead to adaptation, disease, or eventual extinction if left unchecked.

Microbes are known to adopt highly mutable states that would normally be viewed as deleterious in multicellular organisms. Studies have found that microorganisms with defects in pathways associated with genomic integrity, such as those involved in chromosome stability, DNA mismatch repair, DNA damage repair, and cell cycle checkpoints associated with recognizing DNA damage, accelerate adaptation to environmental stressors^1–3^. These defects can be beneficial in the short term, yet deleterious in the long term as mutations continue to accumulate. Defects in DNA mismatch repair resulting in increased mutation rates have been reported in fungi, including the model yeast *Saccharomyces cerevisiae*, the human pathogen *Candida glabrata*, an outbreak strain of *Cryptococcus deuterogattii*, and several clinical isolates of the model basidiomycete human fungal pathogen *Cryptococcus neoformans*^4–11^. Genomic stability in pathogenic *Cryptococcus* species is also significantly affected by karyotypic changes and transposable elements, both of which can mediate antifungal drug resistance^12–16^.

*Cryptococcus* is an environmentally ubiquitous haploid basidiomycete and facultative human pathogen^17^. Approximately 95% of cryptococcal infections are attributable to the serotype A group, *C. neoformans* var. *grubii*, now known as *C. neoformans*, which is divided into four lineages: VNI, VNII, VNBI, and VNBII^18–20^. This species primarily infects immunocompromised individuals and accounts for ∼15% of HIV/AIDS-related deaths^21^. The threat of cryptococcal infections is exacerbated because of the limited antifungal drug arsenal. Amphotericin B, a fungicidal polyene, is often used in combination with 5-flucytosine (5-FC), an antimetabolite, as a first-line treatment strategy for cryptococcal infections^22, 23^. Unfortunately, amphotericin B and 5-FC have undesirable side effects, and 5-FC monotherapy frequently leads to resistance^24–27^. Fluconazole is used to treat asymptomatic patients with isolated cryptococcal antigenemia, those with disease limited to lung nodules or central nervous system infections after clearance of cerebrospinal fluid cultures, or for chronic maintenance therapy^22^. However, *C. neoformans* frequently develops resistance to fluconazole via aneuploidy, particularly Chromosome 1 disomy, or mutations in the sterol biosynthesis pathway, contributing to recurrent infections^12, 28–30^. The limited number of drugs available to treat cryptococcosis, prevalence of resistance and recurrent infections, and difficulty in developing novel antifungal therapies combine to make *C. neoformans* drug resistance an important clinical problem.

Transposons in the C. *neoformans* H99 reference strain and the sister species *Cryptococcus deneoformans* JEC21 reference strain have been characterized^14, 31, 32^. Their genomes encode many retrotransposons both with and without long-terminal repeats, known as LTR retrotransposons and non-LTR retrotransposons, respectively. These retrotransposons move via a copy-and-paste mechanism, allowing them to proliferate throughout the genome if unchecked. The most well-characterized *Cryptococcus* LTR-retrotransposons are Tcn1 through Tcn6, which primarily localize in centromeres^33^. The *C. deneoformans* JEC21 genome also encodes three DNA transposons (T1, T2, and T3), as well as ∼25 copies of the non-LTR retrotransposon Cnl1 (*C. neoformans* LINE-1), which is thought to associate with telomeric repeat sequences^14, 15, 34^. In the *C. neoformans* H99 genome, there are no full-length copies of Cnl1 or DNA transposons^14^.

Studies have illustrated that transposon silencing in *Cryptococcus* is governed by RNA interference (RNAi) through three primary lines of evidence: 1) small interfering RNAs map predominantly to transposons, 2) RNAi mutants show increased transposon expression, and 3) spliceosomes stall on transposon transcripts at an unusually high rate, triggering RNAi^14, 35–39^. Other mechanisms thought to regulate *Cryptococcus* transposons include 5-methylcytosine DNA methylation^33, 40, 41^ and heterochromatic marks^42^. Interestingly, the outbreak species *C. deuterogattii* is RNAi deficient due to the truncation or loss of many genes encoding RNAi components^37^. This loss of RNAi is associated with loss of all functional transposable elements, consequently shorter centromeres, and higher rates of intron retention^33, 43^.

Here, we identified two clinical, hypermutator *C. neoformans* isolates with significantly increased mutation rates on media containing the antifungals rapamycin and FK506. The majority of drug resistance in these two strains was mediated by Cnl1 transposon insertions. Genetic backcrossing, quantitative trait loci mapping, and CRISPR-mediated gene editing confirmed that a nonsense mutation in the RNAi component *ZNF3*, resulting in RNAi loss, is the cause of hypermutation in these strains. Small RNA sequencing confirmed the role of Znf3 in silencing Cn1l, and whole-genome sequencing revealed both hypermutator genomes encode >800 Cnl1 copies or fragments. This is the first time full-length copies of Cnl1 have been identified in *C. neoformans*, and the massive Cnl1 burden in these hypermutators is substantially higher than previously observed in any other *Cryptococcus* strain. Our results demonstrate the hypermutator phenotype described here is attributable to loss of RNAi, allowing rampant transposition of Cnl1. These transposition events lead to Cnl1 accumulation at subtelomeres and movement to novel genomic locations, which can result in drug resistance.

## Results

### Identification of two clinical, hypermutator *C. neoformans* isolates

To identify natural isolates of *C. neoformans* with increased mutation rates, we utilized a collection of 387 *C. neoformans* strains from all lineages (VNI, VNII, VNBI, VNBII), including geographically diverse clinical and environmental isolates of both mating types. For each isolate in this collection, whole-genome sequencing and phylogenetic relationships are available^20^.

Isolates were qualitatively screened for increased mutation rates in a relatively high-throughput manner on medium containing either 5-fluorocytosine (5-FC) or a combination of FK506 and rapamycin (immunosuppressants that bind FKBP12 to form complexes that inhibit activity of calcineurin and TOR, respectively) by evaluating their ability to generate resistant colonies (Figure 1A)^44–47^. For this initial screen, 5-FC was chosen due to its clinical relevance and because genes in which mutations can cause resistance are known^48, 49^. Rapamycin and FK506 were utilized because 1) mutations in only a single gene can mediate resistance to both drugs, and 2) our extensive experience with these drugs, including studies on targets and mechanism of action^50^. Strains that produced more spontaneously resistant colonies on average than the *C. neoformans* H99 reference strain were categorized as hypermutator candidates. We screened 186 strains and identified 36 hypermutator candidates (Table S1). All but one of fourteen (93%) environmental isolates screened were hypermutator candidates (compared to only 14% (23/169) of clinical isolates, *p*-value = 1.1×10^-9^, one-sided Fisher’s exact test). Two previously identified hypermutator strains with mismatch repair defects, C23 and C45, were identified as hypermutator candidates^9^. We chose to focus on two clinical strains, Bt65 and Bt81, that produced the most rapamycin + FK506-resistant (R+F^R^) colonies (Figure 1A). Bt65 and Bt81 are both VNBII

**Figure 1.**
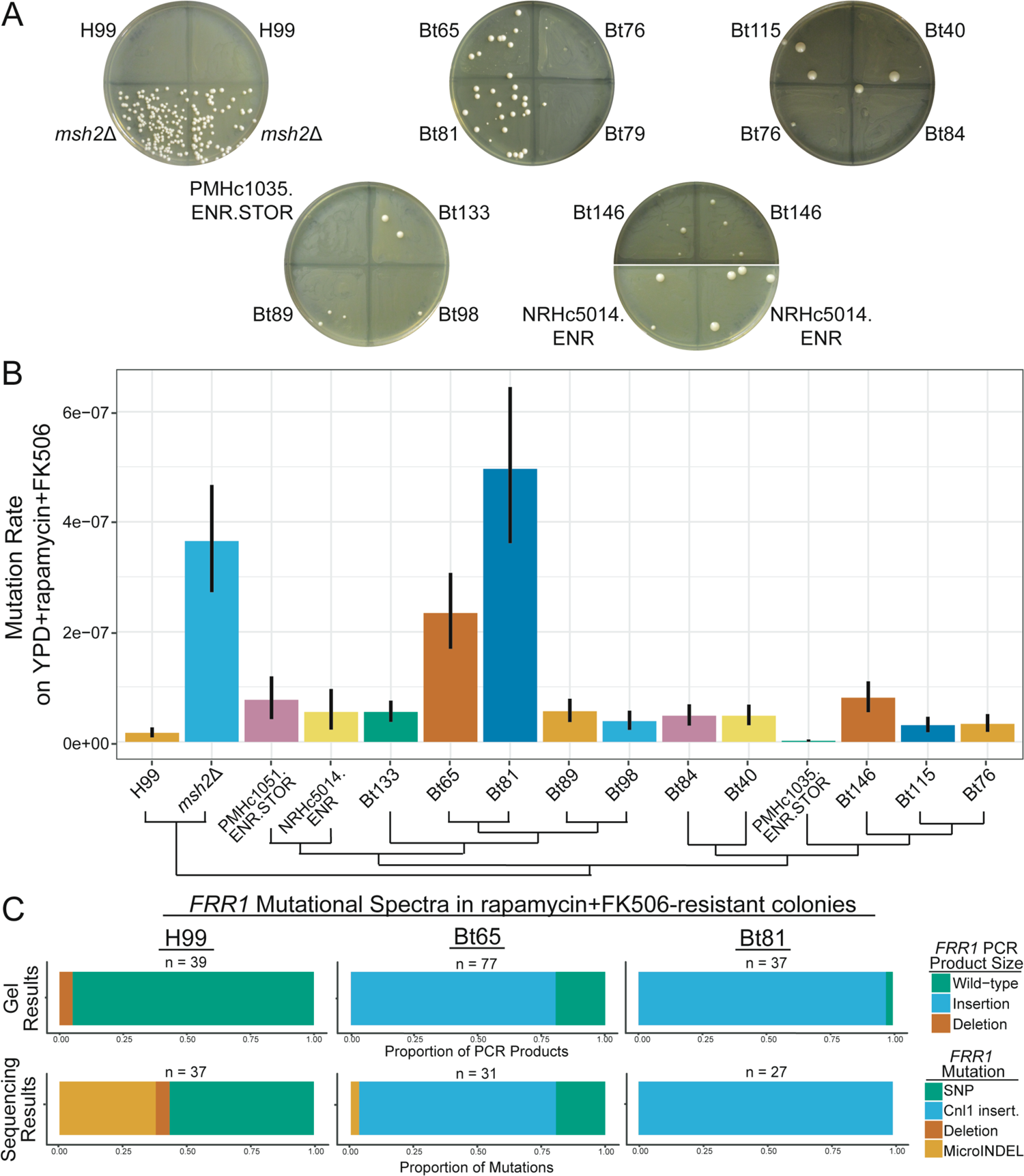
Hypermutation in Bt65 and Bt81 is driven primarily by the insertion of Cnl1 into *FRR1*. **(A)** Generation of spontaneously resistant colonies on YPD + rapamycin + FK506 medium was utilized to identify hypermutator candidates; pictures of representative plates are shown. Strains include the phylogenetically closely related strains involved in fluctuation assay in B as well as positive (*msh2*Δ) and negative (H99) controls. **(B)** Mutation rates of closely related VNBII strains and controls on YPD + rapamycin + FK506. Bars represent the mutation rate and error bars represent 95% confidence intervals; mutation rates represent the number of mutations per cell per generation. Schematic depicts the phylogenetic relationships of all strains included in fluctuation analyses based on Desjardins et al. 2017^20^. Mutational spectra in *FRR1* in YPD + rapamycin + FK506-resistant colonies of H99, Bt65, and Bt81 as characterized by **(C)** gel electrophoresis and Sanger sequencing of *FRR1* PCR products. MicroINDELs are defined as insertions or deletions < 50 bp. All mutations are relative to the appropriate rapamycin + FK506-sensitive parental strain.

*MAT***a** strains from different HIV-positive individuals in Botswana^20, 51^. To quantify the mutation rates of Bt65 and Bt81, we performed fluctuation assays on YPD + rapamycin +FK506 (R+F), YNB + 5-fluoroorotic acid (5-FOA), and YNB + 5-FC media. Both Bt65 and Bt81 produced significantly higher mutation rates on R+F compared to H99 and eleven of the most closely phylogenetically related strains (Figure 1B). On 5-FC, only NRHc5014.ENR and the KN99α *msh2*Δ positive control^52, 53^ had significantly higher mutation rates than H99 (Figure S1A); on 5-FOA (Figure S1B), only KN99α *msh2*Δ produced a significantly higher mutation rate.

A recent study illustrated how incubation at an elevated temperature of 37°C results in increased mutation rates due to transposon mobilization in the closely related species *C*. *deneoformans*^15^. To determine if elevated temperature contributed to hypermutation in Bt65 and Bt81, we concurrently grew these strains as well as wild-type H99 and *msh2*Δ, *ago1*Δ, and *rdp1*Δ deletion mutants overnight at 30°C and 37°C. Fluctuation analysis on R+F medium revealed Bt81 had a significantly lower mutation rate when grown overnight at 37°C compared to 30°C, and Bt65 showed no significant decrease in mutation rate after growth at 37°C; all other strains did not show significant changes (Figure S2). These results suggest that unlike *C. deneoformans*, growth at higher temperature does not contribute to or exacerbate hypermutation in Bt65 and Bt81.

### Characterization of mutation spectra in *C. neoformans* hypermutator strains

We next investigated the types of mutations conferring resistance to the combination of rapamycin and FK506. PCR amplification of the *FRR1* gene (encodes FKBP12, the shared target of rapamycin and FK506 and only gene in which mutations confer resistance to both drugs) followed by gel electrophoresis revealed the expected wild-type product size for all but two (35/37) H99 R+F^R^ colonies; the remaining two produced products smaller than expected, indicative of deletions (Figure S3A). In contrast, large insertions of various sizes were observed in the majority of Bt65 and Bt81 R+F^R^ colonies (62/77 and 36/37, respectively) (Figure 1C and S3B). Only one resistant colony derived from a non-hypermutator strain, Bt84 (1/10 independent colonies), had an insertion in *FRR1*. No insertions in *FRR1* were observed in any other closely related or control strains.

We subsequently sequenced *FRR1* in H99, Bt65, Bt81, and Bt84 R+F^R^ colonies to determine the genetic changes responsible for the varying PCR product sizes (Figure 1C). In 37 H99 R+F^R^ colonies, SNPs in *FRR1* were largely responsible for resistance (57%, 21/37 colonies), while resistance in the remaining colonies was attributable to small (≤ 50 bp) insertions/deletions (microINDELs; 38%, 14/37) or large deletions (5%, 2/37). Conversely, in the hypermutator Bt65, insertions of the non-LTR retrotransposon Cnl1 were responsible for the majority of rapamycin + FK506 resistance (77.4%, 24/31). Rapamycin + FK506 resistance in the remaining Bt65 colonies was either due to SNPs (19.4%, 6/31) or microINDELs (3.2%, 1/31). In all sequenced PCR products from Bt81 R+F^R^ colonies, Cnl1 insertions were responsible for resistance (27/27 colonies). Cnl1 insertions in Bt65 and Bt81 ranged from 54 to ∼3500 bp, and this range in insertion sizes is a common characteristic of non-LTR retrotransposons. The single *FRR1* insertion observed in Bt84 had no homology with any annotated *Cryptococcus* transposons but was identified as a repetitive element by RepeatMasker and shared minor homology with a Copia-58 BG-I transposon.

5-FC- and 5-FOA-resistant colonies of Bt65, Bt81, and H99 were similarly characterized to determine the sources of resistance to antifungal drugs with different mechanisms of action. Resistance to 5-FOA is conferred by mutations in the *URA3* or *URA5* genes of the uracil biosynthesis pathway^54, 55^. Among the H99, Bt65, and Bt81 5-FOA^R^ colonies sequenced, mutations were only identified in *URA5*. In almost all colonies, resistance was conferred by SNPs or INDELs, and only one Cnl1 insertion was identified in a Bt81 5-FOA^R^ colony (Figure S3C). We also PCR amplified genes in which mutations are known to confer resistance to 5-FC, including *FUR1* and *UXS1*^48^. Of the 5-FC^R^ isolates analyzed from H99, Bt65, and Bt81, PCR and sequencing revealed Cnl1 insertions in *FUR1* in two 5-FC^R^ Bt65 isolates and five 5-FC^R^ Bt81 isolates (Figure S3D). Cnl1 insertions into UXS1 were also observed to confer 5-FC resistance in one 5-FC^R^ isolate from both Bt65 and Bt81 (Figure S3E).

Analysis of the Cnl1 insertions observed to confer resistance to R+F, 5-FC, and 5-FOA revealed Cnl1 preferentially inserts at guanine- and cytosine-rich regions of target genes, a known property of this element^32^. Target-site duplication sequences flanking Cnl1 insertions were not present in many instances, but when present, ranged from 1 to 12 bp in length. Cnl1 insertions ranged greatly in size, from 25-bp fragments to full-length Cnl1 copies (3,494 bp). The smallest Cnl1 insertion (25 bp) was followed immediately by a 59-bp deletion in *FRR1*. Of the 51 characterized Cnl1 *FRR1* insertions, 27 were in the 5’ UTR (26/27 were oriented 5’ to 3’, the same orientation as *FRR1* transcription), 23 were in exons (7 oriented 5’ to 3’, 16 oriented 3’ to 5’), and one insertion was in an intron of *FRR1* in the 3’ to 5’ orientation, potentially disrupting splicing or transcription.

### QTL mapping identifies loci that significantly contribute to the hypermutator phenotype

To determine the genetic cause of the hypermutator phenotype and rampant transposition in Bt65 and Bt81 and to determine the genetic consequences of this phenotype, we conducted quantitative trait locus (QTL) mapping. For this purpose, a total of 165 basidiospores were dissected from a genetic cross between Bt65 *MAT***a** and an H99 *crg1*Δ *MAT*α mutant with an enhanced mating phenotype^56^, and 47 F_1_ progeny germinated (28%).

Twenty-eight Bt65**a** x H99α F_1_ progeny were selected for fluctuation analysis and whole-genome sequencing. Alignment of the paired-read Illumina sequencing data from the 28 F_1_ progeny identified 215,411 bi-allelic SNPs that were utilized for QTL mapping. For 24 of the segregants as well as for the Bt65 and H99 *crg1*Δ parental strains, the mutation rate on rapamycin + FK506 medium served as the phenotype for association tests (Figure 2A). Across the 14 chromosomes and bi-allelic SNP sites, two QTL with large effect (heritability = 64%) were identified at approximately 919-1,120 kb on Chromosome 3 (Chr3) and 987-1,193 kb on Chr11 (Figure 2B and S4). Analysis of these QTLs revealed that the SNPs in each QTL were co-segregating and that they shared the same distributions of phenotype scores (Figure S4 and S5). The borders of the QTL spanning Chr3 and Chr11 were determined by calculating 95% confidence intervals and examining recombination break points along each chromosome. Interestingly, these two QTLs span the chromosomal translocation between Chr3 and Chr11 that is unique to H99 (Figure S4, S5, S6). We thus treated these QTLs as the same QTL for subsequent analysis.

**Figure 2.**
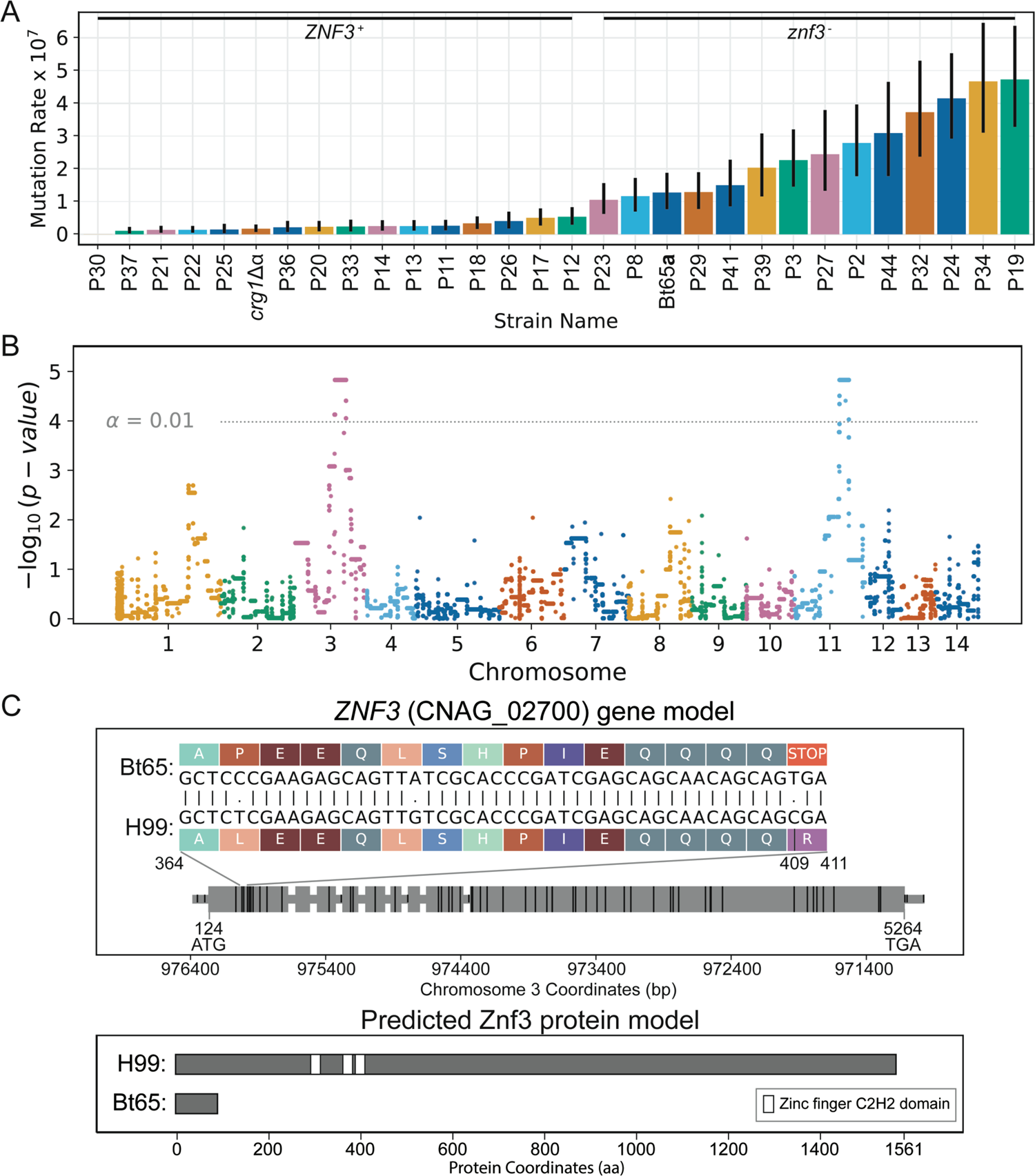
QTL analysis of hypermutator phenotype. (**A**) Quantification of mutation rates on YPD + rapamycin + FK506 medium – sorted smallest to largest, left to right – for F1 progeny and parental strains, H99 *crg1*Δ and Bt65. Inheritance of the Bt65 *znf3* allele or H99 *crg1*Δ *ZNF3* allele in F1 progeny is indicated above mutation rates. Colored bar plots and vertical black lines depict the mean mutation rate and associated 95% confidence intervals (CI) per segregant. Mutation rates represent the number of mutations per cell per generation. (**B**) Manhattan plot showing the strength in association (y-axis) between bi-allelic SNPs and hypermutator phenotype, across the 14 chromosomes (x-axis). Colors separate SNPs across chromosomes. The permutation-based significance threshold (α = 0.01) is depicted with a horizontal dashed line. (**C**) Predicted *ZNF3* gene and Znf3 protein models in H99 and Bt65. A grey horizontal bar depicts the gene body in the upper panel, and larger grey rectangles represent exons; the gene is depicted 5’ to 3’ and is 5417 nt in length. The locations of SNPs differing between Bt65 and H99 are shown by vertical black rungs along the gene model. Amino acids specified by mRNA codons in the indicated region of *ZNF3* exon 1 (nucleotides 364 to 411) are shown for H99 and Bt65 to illustrate the effect of the C to T mutation (nucleotide 409) predicted to cause a nonsense mutation in Bt65. The bottom panel depicts the predicted impact of the nonsense mutation on the Znf3 protein in Bt65. White rectangles along the protein schematic depict the three C2H2-type zinc finger domains of Znf3.

Within the QTL there are a total of 108 and 85 genes along Chr3 and Chr11, respectively, and for 82 and 77 of these genes (respectively), the published annotation and SNP data was used to characterize differences in predicted protein sequence and expected protein lengths between the H99 and Bt65 parental strains (Figure S4 and Table S2). Among these, 71 and 60 genes along Chr3 and Chr11, respectively, have at least one predicted nonsynonymous change in protein sequence, seven of which harbor a predicted nonsense (i.e. stop-gain) or stop-loss mutation. One of these genes is *ZNF3* (CNAG_02700), which encodes a C2H2 type zinc finger protein with three zinc finger domains. Znf3 was previously identified as an RNAi silencing component that localizes to P-bodies and whose mutation results in increased expression of transposable elements^37, 38^. *ZNF3* is located on Chr3 and has a SNP – C to T – within the first exon in the Bt65 genetic background, which is predicted to cause a nonsense mutation, severely truncating Znf3 from 1,561 amino acids to only 96 amino acids (Figure 2C). In addition, this nonsense mutation may also result in nonsense-mediated mRNA decay of the mutant *znf3* mRNA. Based on the publicly available whole-genome sequencing of all isolates in the SDC, the *znf3* nonsense mutation in exon 1 is unique to Bt65 and Bt81 and not present in any other strain. Another gene of known function within the QTL encodes a long-chain acyl-CoA synthetase (CNAG_01836, Chr11) and a SNP – G to A – within the last exon of this gene is predicted to cause an early nonsense mutation in the Bt65 background (Figure S7). Given the dramatic difference in the predicted protein length of *ZNF3* between the H99 and Bt65 parental alleles (relative to other genes in this QTL with predicted stop-loss or nonsense mutations), and previous studies demonstrating the role of Znf3 in RNAi and transposon silencing, we hypothesized *ZNF3* could be the quantitative trait gene (QTG) and the SNP leading to the predicted stop gain in the first exon could be the quantitative trait nucleotide (QTN) underlying the hypermutation phenotype^37, 38^.

### Few Bt81 F_1_ progeny display a hypermutator phenotype

Forty-two F_1_ progeny were also derived from a genetic cross between the other hypermutator strain, Bt81**a,** and H99α *crg1*Δ. The *ZNF3* alleles of all Bt81 F_1_ progeny were sequenced to determine whether each progeny had inherited the non-functional Bt81 *znf3* allele or the functional H99 *ZNF3* allele. Of the 42 F_1_ progeny, only four inherited the mutant *znf3* allele from Bt81, a significantly lower number than expected based on Mendelian inheritance patterns (chi-square test, *p*-value < 0.01). The four progeny with non-functional Bt81 *znf3* alleles had the highest mutation rates of 18 F_1_ progeny that were analyzed (Figure S8). However, three of the *znf3* progeny had mutation rates that were not significantly higher than those of the Bt81 *ZNF3* progeny and were also not as high as would be expected based on results from the Bt65**a** x H99α F_1_ progeny (Figure 2A).

### Cnl1 elements are organized into subtelomeric arrays in hypermutator genomes

For all strains in the SDC, including Bt65 and Bt81, only short-read whole-genome sequencing data was available^20^. Because of the known difficulties in assembling repetitive elements, such as Cnl1, with short-read sequencing data, we conducted long-read whole-genome sequencing with the Oxford Nanopore Technologies MinION to generate more complete assemblies for Bt65, Bt81, and two of the most closely phylogenetically related non-hypermutator strains, Bt89 and Bt133. With long-read sequencing data, we assembled chromosome-level genomes for all four strains. In the assemblies, we observed the known chromosomal translocation between Chr3 and Chr11 unique to H99^57^ and identified a translocation between Chr1 and Chr13 unique to Bt65 and Bt81 (Figure S6). These two gross chromosomal rearrangements explain the relatively low germination frequency (28%) of Bt65**a** x H99⍺ F_1_ progeny because each translocation should decrease germination by ∼50%.

Analysis of the genomes of Bt65 and Bt81 revealed large arrays of the Cnl1 transposon at all but one end of each of the 14 linear chromosomes (27/28 subtelomeric regions in Bt65 and 28/28 in Bt81) (Figure 3A, 3B). The assembled Cnl1 arrays (defined as ≥ 2 Cnl1 copies) in Bt65 and Bt81 range from 5 kb to 80 kb in length. These highly repetitive arrays made it difficult and, in some instances, impossible to confidently assemble telomeric repeat sequences at the ends of each Bt65 and Bt81 chromosome. Using manual telomere extension via read mapping, we were able to identify telomere repeats at only 20 chromosome ends in Bt65 and 13 in Bt81. In contrast to Bt65 and Bt81, genome assemblies for Bt89 and Bt133 were assembled with telomere repeats on all 28 chromosome ends without any manual extension (Figure S6). In these assemblies, some telomeres had no copies of Cnl1 while others had Cnl1 arrays up to 30 kb in length (Figure S9).

**Figure 3.**
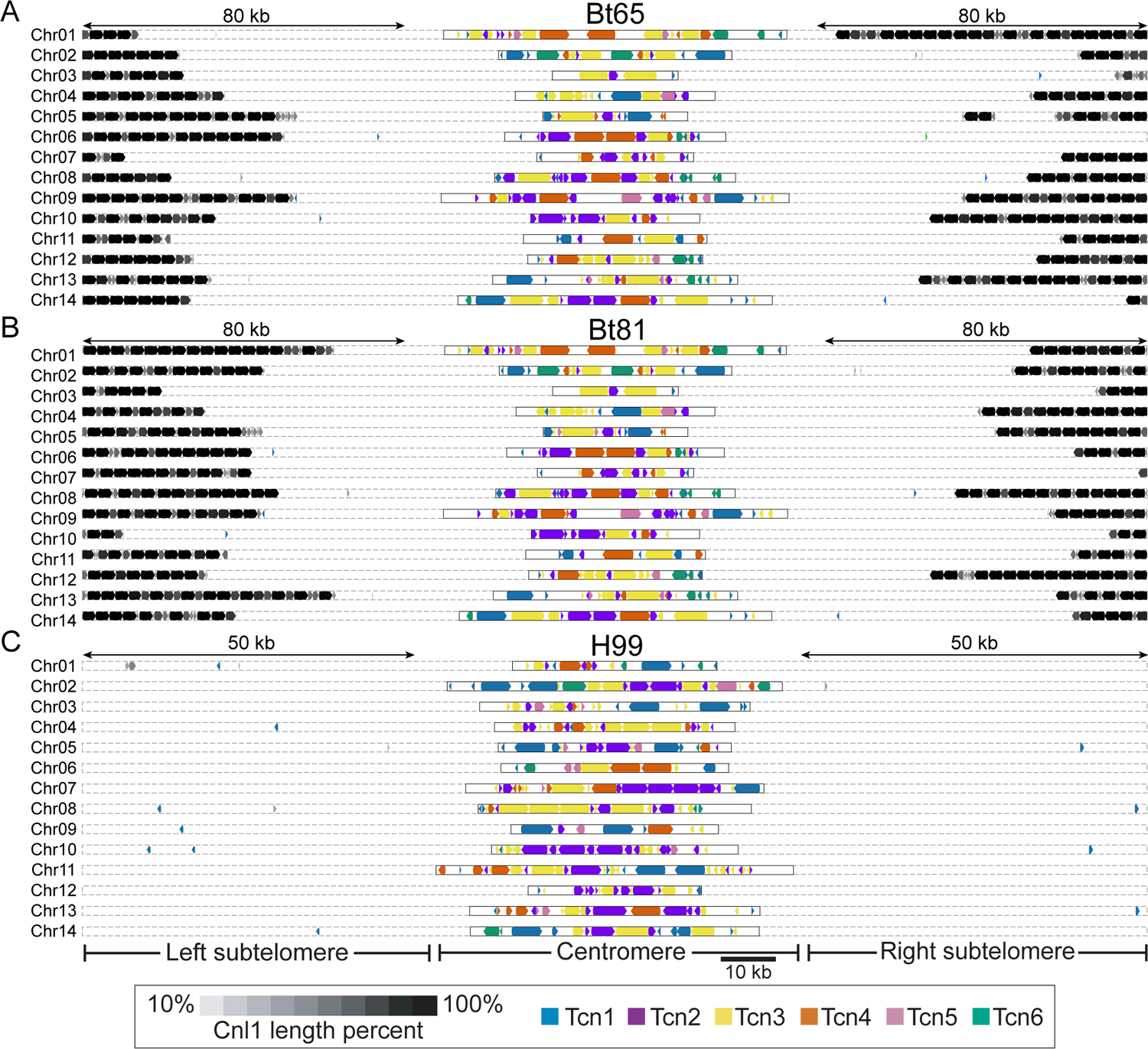
Retrotransposon content in the genomes of H99, Bt65, and Bt81. Distributions of the Tcn1 through Tcn6 LTR-retrotransposons and the Cnl1 non-LTR retrotransposon in subtelomeric and centromeric regions of **(A)** Bt65, **(B)** Bt81, and **(C)** H99 genomes depicted in Figure S6. In Bt65 and Bt81, 80 kb of subtelomeric regions are displayed, and 50 kb subtelomeric regions are displayed for H99 to show the full distribution of subtelomeric Cnl1 elements. Subtelomeric arrays of Cnl1 are depicted at the end of each chromosome in Bt65 and Bt81, while only 7 Cnl1 elements are localized subtelomerically in H99. Shading corresponds to fragments of the Cnl1 elements, and gene arrowheads indicate the direction of transcription for all retrotransposons.

Further analysis revealed the Bt65 genome harbors at least 414 fragments of Cnl1, including 105 full-length copies, while the Bt81 genome appears to encode even more Cnl1 elements, with at least 449 fragments, including 147 full-length copies (Table 1). It is important to note that due to incomplete ends for most chromosomes, Bt81 and Bt65 likely encode additional, unassembled Cnl1 copies. The presence of long Cnl1 arrays in Bt65 and Bt81 was surprising because the H99 reference strain encodes only 22 fragments of Cnl1 and no full-length copies; *C. neoformans* was therefore not thought to harbor functional Cnl1 elements (Figure 3C) (Table 1).

**Table 1.** Cnl1 burden in H99, hypermutator strains, related non-hypermutator strains, and six Bt65 x H99 *crg1*Δα F_1_ progeny based on Nanopore sequencing data.

| Strain | Hypermutator Status | Total Cnl1 burden<br>(>50 bp) | Full-length Cnl1 copies<br>(>99% in length) |
| --- | --- | --- | --- |
| H99 | Non-hypermutator | 22 | 0 |
| Bt65 | Hypermutator | 414 | 105 |
| Bt81 | Hypermutator | 449 | 147 |
| Bt89 | Non-hypermutator | 261 | 24 |
| Bt133 | Non-hypermutator | 246 | 23 |
| Progeny 2 | Hypermutator | 212 | 30 |
| Progeny 8 | Hypermutator | 172 | 40 |
| Progeny 14 | Non-hypermutator | 296 | 68 |
| Progeny 18 | Non-hypermutator | 321 | 88 |
| Progeny 20 | Non-hypermutator | 425 | 136 |
| Progeny 34 | Hypermutator | 187 | 41 |

Apart from the subtelomeres, retrotransposons in *Cryptococcus* are also enriched at centromeres, specifically the LTR retrotransposons Tcn1-Tcn6^31, 33, 57^. The changes in Cnl1transposon content in Bt65 and Bt81 along with a previous study establishing a link between RNAi loss and centromere length^33^ motivated us to characterize the centromeres in Bt65, Bt81, Bt89, and Bt133. Analysis revealed shorter centromeres on average in these isolates compared to H99. However, this difference did not reach statistical significance (ANOVA, *p*-value = 0.153, Figure S10 and Table S3). Many centromeres in the assessed isolates had undergone numerous rearrangements and inversions relative to one another (Figure S11). Centromeric alterations have also previously been observed in *C. neoformans* genetic deletion mutants lacking the canonical RNAi components Ago1 and Rdp1^33^. Combined, these analyses suggest that while Cnl1 is more abundant in Bt65, Bt81, Bt89, and Bt133, other retrotransposons are not substantially increased in number compared to H99 (Figure 3 and S9).

### Characterization of H99 *crg1*Δ x Bt65 F_1_ progeny genomes reveals invasion of Cnl1 elements into naïve telomeres

Expression of transposable elements, including Cnl1, is upregulated during sexual reproduction in *C. neoformans*^35, 37, 39^. To investigate how increased expression of Cnl1 during mating impacts the genome, six progeny utilized for QTL mapping were also selected for long-read whole-genome sequencing: three *znf3* hypermutator progeny (P2, P8, and P34) and three *ZNF3* non-hypermutator progeny (P14, P18, and P20). Long-read sequencing identified recombination points across the progeny genomes, providing information on which regions were inherited from either parent and confirming these were F_1_ genetic recombinants (Figure 4).

**Figure 4.**
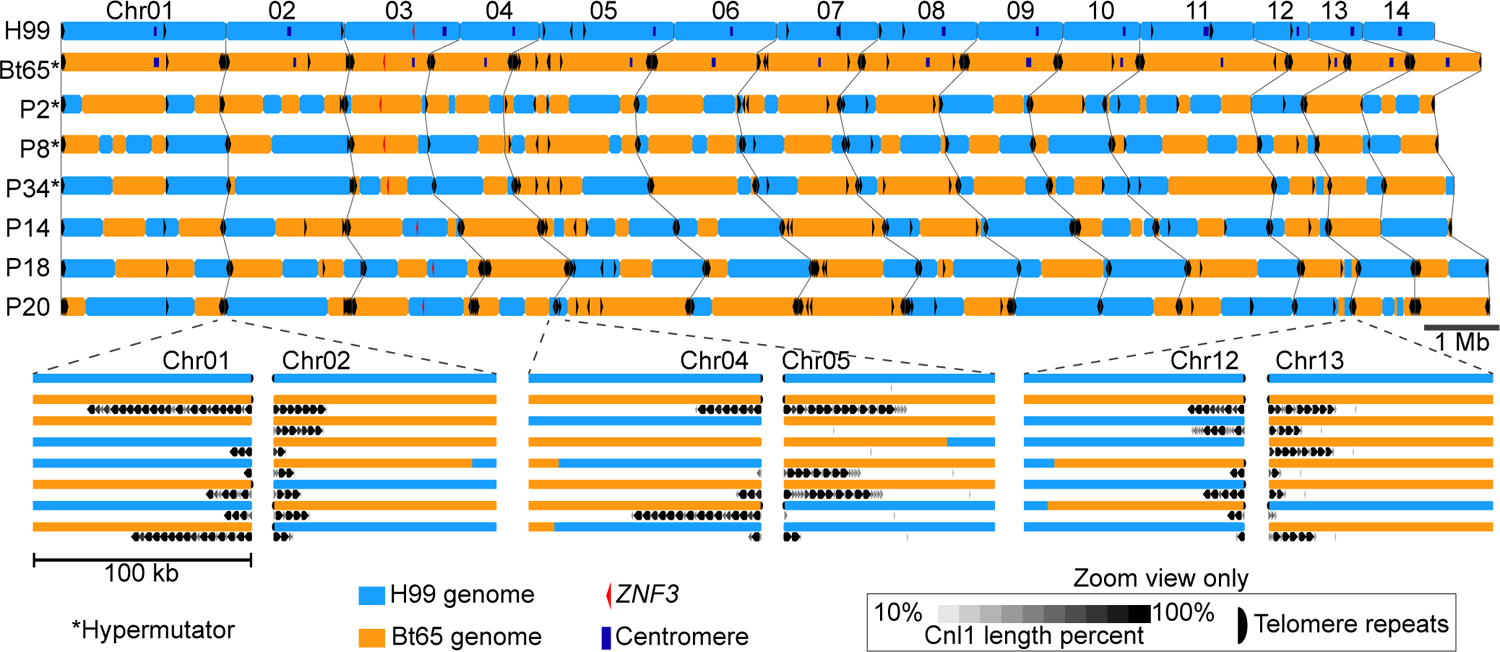
Genetic recombination sites and Cnl1 distribution in Bt65 x H99 F1 progeny. Recombination sites along each of the 14 chromosomes for the six Bt65**a** x H99α F1 progeny for which long-read whole-genome sequencing was conducted. Genomic loci depicted in blue were inherited from the H99 parent, and orange genomic loci were inherited from the Bt65 parent. Cnl1 elements throughout the F1 progeny and parental genomes are indicated by black arrowheads in the upper panel. Centromeres are indicated by dark blue boxes in only the parental genomes. Hypermutator F1 progeny are indicated with asterisks, and the *ZNF3* locus is indicated in each strain with a red arrowhead. Regions enlarged below illustrate Cnl1 subtelomeric arrays on several chromosomes and depict examples of Cnl1 array expansion (e.g. Chromosome 4, P18), contraction (e.g. Chromosome 1, P14), and invasion of naïve H99 subtelomeres (e.g. Chromosome 1, P8). Telomeric repeat sequences are indicated by black half circles only in the enlarged panels.

Surprisingly, the genome assemblies for the three *ZNF3* non-hypermutator progeny appear to encode more full-length Cnl1 elements and fragments than the three *znf3* progeny. However, of the three *znf3* progeny, telomeric repeat sequences were only identified at the end of two chromosomes (2/84 telomeric ends across three progeny, 2%) (Figure S12). This is in contrast to the 31 telomeres accurately assembled across the three *ZNF3* progeny (31/84, 37%) (Figure S12). The smaller number of telomeres identified in *znf3* progeny suggests more Cnl1 elements may not have been accurately included, similar to the assemblies for Bt65 and Bt81. Therefore, the Cnl1 content in Table 1 might not accurately capture the entire Cnl1 burden in these strains.

A previous study found *ZNF3* to be a haploinsufficient gene because no progeny from a *ZNF3* x *znf3*Δ cross showed evidence of sex-induced RNAi-mediated silencing^37^. This haploinsufficiency allowed us to analyze Cnl1 dynamics in both hypermutator and non-hypermutator progeny. Nearly all Cnl1 arrays in the progeny showed signs of expansion and contraction relative to Bt65, suggesting these elements are highly mobile during mating, undergoing high levels of recombination, or both.Combined analysis of subtelomeric inheritance patterns and Cnl1 arrays revealed Cnl1 is capable of invading naïve subtelomeres inherited from H99, i.e. regions previously devoid of Cnl1 (Figure 4, Figure S12). In the three *znf3* progeny, 65% (28/43) of naïve subtelomeric regions inherited from H99 acquired Cnl1 copies, and arrays in many cases. In the three *ZNF3* progeny, 81% (35/44) of naïve subtelomeres from H99 now had Cnl1 elements. Overall, both *ZNF3* and *znf3* progeny inherited roughly equivalent numbers of subtelomeric regions from either parent and Cnl1 invaded a majority of the naïve H99 telomeres.

### Replacement of the *ZNF3* nonsense mutation significantly lowers the mutation rate and restores siRNA production

The evidence presented thus far suggested the nonsense mutation in *ZNF3* unique to Bt65 and Bt81 is responsible for the hypermutation phenotype, possibly due to compromised RNAi silencing of Cnl1. To test this hypothesis, we used CRISPR-mediated gene editing to restore the functional *ZNF3* allele in Bt65. Gene editing was achieved with the TRACE system^58^ and utilization of a functional *ZNF3* allele from the closely related strain Bt133, such that only the SNP causing the nonsense mutation would be changed to the wild-type nucleotide (found in H99 and all isolates except Bt65 and Bt81)^58^. Following transformation and selection, we identified two independent Bt65 transformants that had successfully integrated a single copy of the Bt133 *ZNF3* allele at the endogenous *znf3* locus, Bt65+*ZNF3*-*1* and Bt65+*ZNF3*-*2*. These transformants were subjected to fluctuation analysis to determine if reverting the *znf3* nonsense mutation restored the mutation rate to a wild-type level. On R+F medium, both transformants had significantly lower mutation rates than Bt65, similar to H99 and Bt133 (Figure 5A).

**Figure 5.**
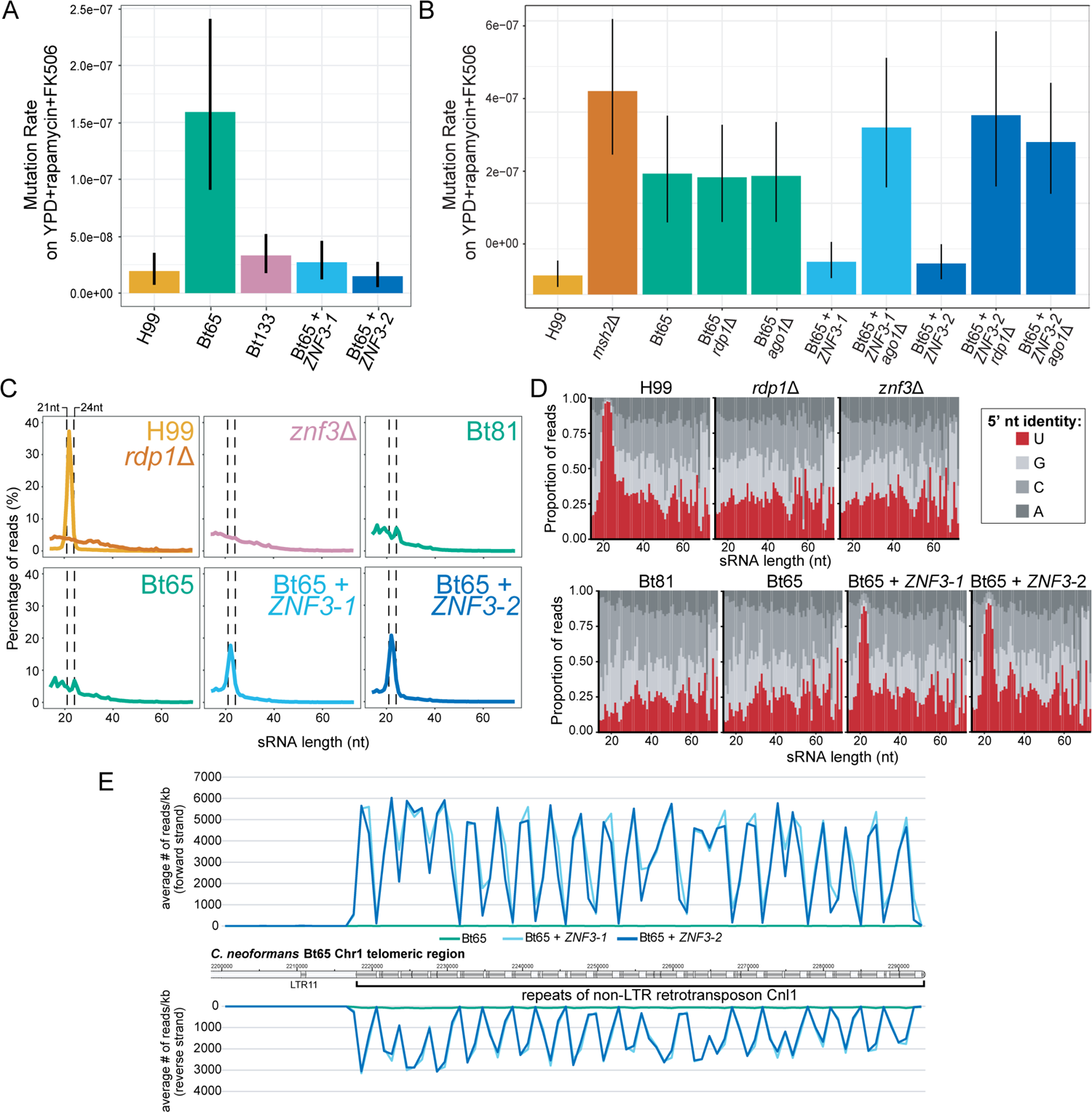
*ZNF3* complementation in Bt65 significantly reduces mutation rates and restores siRNA production. Mutation rates of **(A)** the two independent *ZNF3* complementation mutants, Bt65+*ZNF3*-*1* and Bt65+*ZNF3*-*2*, as well as control strains, and **(B)** *ago1*Δ and *rdp1*Δ deletion mutants in the Bt65, Bt65+*ZNF3*-*1* and Bt65+*ZNF3*-*2* genetic backgrounds and controls on YPD+rapamycin+FK506 medium. Bars represent mutation rate (number of mutations per cell per generation) and error bars represent 95% confidence intervals. **(C)** Size distributions of sRNA reads from each indicated strain. Dashed vertical lines indicate the 21 to 24 nucleotide size range, the characteristic sizes of siRNAs produced by the RNAi pathway. **(D)** Proportion of sRNA reads (y-axis) with the indicated 5’ nucleotide identity (color of stacked bar) at each sRNA read size (x-axis). siRNAs produced by the RNAi pathway characteristically have a 5’ uracil nucleotide. **(E)** Quantification of sense and antisense sRNAs from Bt65, Bt65 + *ZNF3*-*1*, and Bt65 + *ZNF3*-*2* aligning to an array of subtelomeric Cnl1 elements on Chromosome 1 of Bt65. Transposable elements along the chromosome are indicated by dark grey boxes, while intergenic regions are light grey.

Because *ZNF3* is required for sexual reproduction in *C. deneoformans*, although not essential for sexual development in *C. neoformans*^37, 38^, we sought to confirm the phenotypes related to *ZNF3* loss and restoration were due specifically to RNAi-mediated silencing. Two canonical RNAi components, *AGO1* and *RDP1*, were therefore genetically deleted in Bt65 and the two Bt65+*ZNF3* strains. The mutation rates of these strains were subsequently measured on R+F medium (Figure 5B). Deletion of *AGO1* or *RDP1* in Bt65 did not affect the mutation rate on R+F. Conversely, deletion of *AGO1* in Bt65+*ZNF3*-*1* and -*2* and deletion of *RDP1* in Bt65+*ZNF3*-*2* led to significantly increased mutation rates, illustrating that it is indeed the loss of the role of Znf3 in RNAi that results in the hypermutator phenotype in Bt65. This finding also provides evidence that all other RNAi components have been largely maintained despite loss of RNAi activity in Bt65, in accord with genome inspection that failed to reveal loss-of-function muations in other RNAi components (*CPR2*, *DCR1*, *QIP1*, *RDP1*, *GWC1*, *FZC28*, *GWO1*, *AGO1*, *DCR2*, *DBR1, RDE1, RDE3, RDE4, RDE5, RPA32, SRR1, RDE2*).

We next sequenced the sRNA repertoires of the Bt65+*ZNF3* transformants as well as Bt65, Bt81, and an H99 *znf3*Δ mutant; H99 and an H99 *rdp1*Δ mutant served controls. Analysis of the size distribution of sRNAs showed that Bt65, Bt81, and H99 *znf3*Δ displayed profiles similar to that of the *rdp1*Δ mutant, lacking the characteristic 21-24 nt siRNA peak (Figure 5C). Analysis of sRNAs also revealed that *ZNF3* complementation in Bt65 restored the 21-24 nt sRNA peak. We also characterized the 5’ nucleotide identity of sRNAs and found only H99 and the two Bt65+*ZNF3* transformants had a peak of 21-24 nt sRNAs with a predominance of 5’ U identity, another characteristic of siRNAs (Figure 5D).

To determine how restoration of *ZNF3* impacted silencing of Cnl1, we quantified sRNAs aligning to Cnl1 elements across the Bt65 genome (Table S4). Relative to Bt65, normalized expression of sRNAs corresponding to Cnl1 were increased 5.6-fold in H99, 10.6-fold in Bt65+*ZNF3*-*1*, and 12.8-fold in Bt65+*ZNF3*-*2* (Table S4). To illustrate the marked difference in Cnl1 sRNAs in the Bt65+*ZNF3* transformants compared to Bt65, sRNAs were plotted along a Cnl1 array on Chr1 in Bt65 (Figure 5E). These results show that changing the single nucleotide responsible for the nonsense mutation in *ZNF3* back to the wild-type nucleotide found in closely related strains as well as H99 was able to successfully restore production of siRNAs in Bt65.

## Discussion

Transposable element mobilization can alter gene expression, gene function, and even genomic stability. In this study, we identified forty *C. neoformans* hypermutator candidates, which were particularly enriched for environmental isolates. Environmental niches may possess a more diverse array of stressors than the human host and thus favor hypermutation. Additionally, our methods for identifying hypermutators with rapamycin + FK506 medium could bias results in favor of strains more likely to modulate FKBP12 activity. FKBP12 modulation may be more important in environmental isolates because rapamycin and FK506 are naturally derived compounds produced by the soil-resident bacteria^59, 60^ and both TOR and calcineurin are instrumental for a variety of stress responses^61, 62^.

Two of the strongest hypermutators, clinical isolates Bt65 and Bt81, were found to have a massive accumulation of the non-LTR retrotransposon Cnl1 at subtelomeric loci. Cnl1 elements were capable of inserting into non-subtelomeric regions of the genome, resulting in resistance to diverse antifungal drugs, namely 5-FOA, the clinically relevant drug 5-FC, and the combination of rapamycin and FK506. These findings were unprecedented as *C. neoformans* is thought to be an RNAi-proficient species and to lack full length copies of Cnl1; only the sister species *C. deneoformans* harbors full-length Cnl1 elements capable of mobilization^14, 32, 63^. These findings highlight the importance of intraspecific diversity at both genotypic and phenotypic levels. It is important to note that the fluctuation assays employed to quantify hypermutation only inform us on the mutation rates at a limited number of loci and therefore cannot inform us on the global mutation rate across the genome. However, we were able to demonstrate that Cnl1 could insert into at least four different genes to confer resistance to each type of media assessed (rapamycin+FK506, 5-FOA, and 5-FC). This observation, combined with the apparent dynamic changes occurring in Cnl1 subtelomeric arrays suggest that these strains are undergoing additional mutagenesis not observed in other strains. We posit that 1) the elevated mutation rate observed at the *FRR1* locus, 2) the Cnl1 insertions at additional genomic loci when selected for (*FUR1*, *UXS1*, *URA5*), and 3) the dynamic changes observed in Cnl1 subtelomeric arrays combine to justify qualification of Bt65 and Bt81 as hypermutators.

Following isolation, phenotyping, and genotyping of the Bt81**a** x H99α F_1_ progeny, a significantly smaller number of progeny inherited the Bt81 *znf3* allele than expected, a surprising result given the nearly 1:1 inheritance of *ZNF3* from either parent in Bt65**a** x H99α F_1_ progeny. Based on long-read whole-genome sequencing, Bt81 has a substantially higher burden of Cnl1 than Bt65. Previous studies have shown that expression of transposons, including Cnl1, is significantly upregulated during sexual reproduction in RNAi mutants, like H99 *znf3*Δ^35, 37–39^. The higher Cnl1 burden in Bt81 combined with RNAi-deficiency and transposon upregulation during mating could lead to an increased frequency of deleterious Cnl1 insertions in progeny lacking *znf3*, and thus biased inheritance of *ZNF3*. It is also possible that a higher Cnl1 burden favored the selection of a suppressor mutation in the Bt81 F_1_ progeny. The mutation rates of the progeny inheriting the nonfunctional *znf3* allele also had mutation rates lower than expected, lending further support to the idea that suppressor mutations limiting Cnl1 movement may be segregating or arising during the cross. Additionally, *znf3*Δ mutants are the only *C. neoformans* RNAi mutants in which progeny from unilateral genetic crosses (i.e. crosses in which only one parent lacks *ZNF3*) exhibit complete loss of RNAi-mediated silencing^37^, and thus loss of *ZNF3* results in haploinsufficiency. This haploinsufficiency could also favor gene conversion events resulting in a *ZNF3*^+^/*ZNF3*^+^ homozygote in the diploid formed during mating prior to meiosis. Overall, unequal *znf3* inheritance patterns in Bt81 F_1_ progeny suggest a sufficiently high burden of Cnl1 may be deleterious during sexual reproduction.

In Bt65, QTL mapping and genetic complementation demonstrated the hypermutator phenotype was caused by a single SNP in *ZNF3*. Changing the *znf3* nonsense mutation to the nucleotide found in the reference strain as well as phylogenetically closely related strains lowered the mutation rate to a wild-type level and restored siRNA production, including siRNAs corresponding to Cnl1. Furthermore, deletion of *AGO1* or *RDP1*, two genes encoding canonical RNAi components, in the Bt65+*ZNF3* strains restored the hypermutator phenotype, confirming that the *znf3* nonsense mutation leads to loss of RNAi and results in increased transposition and hypermutation. Although *ZNF3* complementation restored the mutation rate and siRNA production, the identified QTL spanning Chr3 and Chr11 accounted for only 64% of the hypermutator phenotype. This may suggest the existence of additional contributing small-effect loci, although the mapping population used here is underpowered to detect such loci. Variation in Cnl1 subtelomeric arrays across the progeny may also account for the additional genetic loci contributing to the hypermutator phenotype. However, these Cnl1 loci were likely difficult to map in the F_1_ progeny or failed to pass quality criteria and were filtered out during QTL analysis.

Despite having the wild-type nucleotide in the first exon of *ZNF3*, where Bt65 and Bt81 have a nonsense mutation, both Bt89 and Bt133 (two of the most closely related strains) have a substantial accumulation of subtelomeric Cnl1 arrays. One parsimonious explanation for the considerable Cnl1 burden in Bt89 and Bt133 could be that Cnl1 accumulation predated RNAi loss, with subtelomeric arrays undergoing expansion and contraction via recombination as opposed to transposition. Additional modifiers in these genetic backgrounds mitigating the impacts of rampant Cnl1 transposition may have allowed the persistence of the RNAi-deficient Bt65 and Bt81 strains. Alternatively, these isolates may have descended from an ancestral strain that had lost (via mutation/suppression of *ZNF3* or another RNAi component) and subsequently regained RNAi function, possibly through a genetic cross and inheritance of a functional allele. The bias against *znf3* inheritance in Bt81 F_1_ progeny and the lower but still impressive Cnl1 burden in Bt89 and Bt133 potentially illustrate a natural example of how *C. neoformans* genomes have struck a balance in their mutational capability, switching between high mutational capacities during times of RNAi loss, and genomic stability when RNAi is restored.

The expansion and contraction of Cnl1 arrays and ability of Cnl1 to invade naïve subtelomeric regions inherited from H99 in the Bt65 F_1_ progeny was also exceptional. The combination of this observed Cnl1 invasion into naïve telomeres during mating and the fact that Bt65 and Bt81 are both the rare mating type **a** and members of one of the most frequently recombining *C. neoformans* lineages (VNBII)^20^ suggest Cnl1 may spread and proliferate rapidly in environmental strains. The observed Cnl1 subtelomeric dynamics mirror those observed for MoTeR transposons of the fungal plant pathogen *Magnaporthe oryzae*, which were also shown to localize to dynamic subtelomeric arrays^64^. The Cnl1 subtelomeric arrays identified here could also potentially overcome the requirement for telomerase, as in *Drosophila* telomeres, in which the functions of telomerase have been supplanted by a telomeric retrotransposon^65, 66^.

The finding that only a single SNP rendered the RNAi pathway non-functional in Bt65, and that no additional obvious mutations had occurred in other RNAi genes, suggest Bt65 could illustrate the natural consequences of relatively recent RNAi loss. Further characterization of Bt65 (and potentially Bt81) through experimental evolution or gene regulation analyses could shed light on the short-term consequences of RNAi loss at genomic and phenotypic levels. Bt65 could thus serve as an interesting intermediate evolutionary comparator between RNAi-proficient *C. neoformans* isolates and the closely related RNAi-deficient species *C. deuterogattii*^7, 37^. Studying the dynamics of the Cnl1 subtelomeric arrays following passaging would be particularly interesting, and the immediate impacts of RNAi loss on these arrays could be investigated by introducing the *znf3* nonsense mutation into the closely related Bt89 or Bt133 strains that have more limited subtelomeric arrays, or through additional genetic crosses.

Instances of relatively recent loss of RNAi have also been observed in a natural *Caenorhabditis elegans* isolate, which has a large deletion in a RIG-I homolog required for RNAi and was shown to be infected with an RNA virus^67, 68^. Unlike the identified *C. elegans* virus-infected strain and several other RNAi-deficient fungal species, such as *Saccharomyces cerevisiae*, *Ustilago maydis*, and several *Malassezia* species, we were unable to identify a dsRNA virus in either of the *C*. *neoformans* hypermutator strains identified here (see Materials and Methods)^63, 69^. It is possible though, that the hypermutators harbor other types of mycoviruses (e.g. ssRNA) that we were unable to detect or that the mycovirus was cured by common microbiological isolation practices^69^.

The identification of this hypermutator phenotype in natural *C. neoformans* clinical isolates has important implications for antifungal drug resistance and potentially other adaptive consequences. Here, we showed Cnl1 insertion could confer resistance to diverse classes of antifungal drugs, namely 5-FC, 5-FOA, and the combination of rapamycin and FK506. Insertion of Cnl1 into other genes, particularly those in the sterol biosynthesis pathway, could confer resistance to amphotericin B and fluconazole, the only other drugs effective for *C. neoformans* treatment^28, 70^. This mechanism of drug resistance also has interesting implications for a novel antifungal approach that utilizes dsRNA to initiate RNAi silencing in fungal plant pathogens^71^. The effects of Cnl1 insertion at non-coding loci, such as promoters and 3’ untranslated regions, could also impact overall genomic stability or alter gene expression to have important phenotypic implications for virulence, similar to the effects of the *Ac*/*Ds* elements of maize, the first transposable elements discovered^72^. Alterations in gene expression might also confer resistance to drugs for which resistance cannot be gained through loss of function mutations. Even if full resistance isn’t acquired, altered gene expression could contribute to antifungal drug tolerance, like the tolerance observed in *Candida albicans*, which contributes to persistent infections in immunocompetent patients^73, 74^.

At this stage it is difficult to know how selection may act upon Cnl1 transposition and accumulation over time. The subtelomeric arrays in Bt65 and Bt81 may undergo cycles of amplification and recombination-mediated contraction allowing them to exploit Cnl1 mutagenesis when under stress, similar to retrotransposons replication cycles in some plants^75, 76^. The sister species *C. deneoformans* seems capable of applying a similar strategy by mobilizing transposons throughout the genome under heat stress^15^. The study by Gusa and colleagues illustrates an example of an environmental change triggering transposition, while our findings show how a genetic change can allow rampant transposition, demonstrating the diverse mechanisms of mutagenesis and potentially adaptation in pathogenic *Cryptococcus* species.

Maintaining an RNAi-deficient background could also be adaptive in the context of viral infection, as has been shown in yeast harboring the killer virus, which outcompete neighboring uninfected strains, and in mice harboring latent herpesvirus, which are protected from the bacterial pathogens *Listeria monocytogenes* and *Yersinia pestis*^77, 78^. Conversely, the mutational impact of Cnl1 mobilization and hypermutation could be highly deleterious over the long term and therefore may not represent a massive contributor to the rise of drug resistance. Natural selection could either favor reversion to a functional RNAi-pathway through mutation of *znf3* or preserve RNAi loss and eliminate full-length transposable elements, as in *C. deuterogattii*^33^. Future research on the potential for Cnl1 insertion to mediate resistance to amphotericin B and fluconazole, and the impact of hypermutation due to Cnl1 mobilization on *in vivo* drug resistance, adaptive potential, and genomic stability over time will be of great interest from basic science and translational perspecitves.

## Materials and Methods

### Strains and growth

The *C. neoformans* strains described in this study are listed in Table S5. Strains were stored at −80°C in liquid yeast extract peptone dextrose (YPD) supplemented with 15% glycerol. Strains were inoculated on YPD agar plates, grown for three days at 30°C, and maintained at 4°C. Due to the hypermutator phenotypes associated with several of the strains in this study, strains were not maintained on YPD agar plates for routine use for more than two weeks; fresh cells from frozen glycerol stocks were inoculated to YPD agar plates as needed.

### Screening for hypermutator candidates

Assays for the emergence of resistance (papillation assays) were conducted as previously described^53^. In brief, ten independent cultures of each strain were grown overnight at standard laboratory conditions in 5 mL liquid YPD medium. Cultures were then spun down, washed, and concentrated in 2 mL dH_2_O. Each culture was swabbed to a single quadrant of either YPD + 100 ng/mL rapamycin + 1 µg/mL FK506 agar medium or YNB + 100 µg/mL 5-fluorocytosine agar medium (no repeated measurements). YPD + rapamycin + FK506 plates and YNB + 5-fluorocytosine plates were incubated for up to seven days at 37°C and 30°C, respectively. Fisher’s exact probability test was used to determine if the associations between environmental isolates and the hypermutator phenotype was statistically significant using the VassarStats online software (http://vassarstats.net).

### Fluctuation assays

Fluctuation assays were conducted as previously described^53^. Briefly, ten independent overnights of each strain were grown overnight in 5 mL liquid YPD medium at 30°C. Cultures were washed three times and resuspended in dH_2_O. Cells from each culture were then plated to a single plate of the appropriate medium (100 μL 10^-5^ cells on YPD, 100 μL 10^-2^ cells on YNB + 5-FC, and 100 μL undiluted cells on YPD + rapamycin + FK506 and YNB + 5-FOA) (no repeated measurements). Drug concentrations were determined such that no wild-type/sensitive colonies will grow. For the increased temperature fluctuation analysis, strains were grown overnight at either 30°C or 37°C before use in fluctuation assays, as indicated. YPD + rapamycin + FK506 plates were incubated and 37°C; all other media was incubated at 30°C. Following incubation at the appropriate temperature for four days (YPD control medium) or 14 days (drug media), the number of colonies on each plate was determined and subsequently utilized to calculate mutation rates. Mutation rates and 95% confidence intervals were calculated using the FluCalc program which utilizes the Ma-Sandri-Sarkar maximum likelihood estimation (MSS-MLE) equations for calculations and also incorporates plating efficiency into calculations to account for dilutions^79^; no data was excluded in calculations. In Figures 1 and S1, 20 strains were used to calculate mutation rates (*n* = 20), with the exception of strains PMHc1051.ENR.STOR and NRHc5014.ENR, for which *n* = 10. In Figure S8, *n* = 10 strains were used for all mutation rate calculates, except for the control strains, Bt81 and H99 *crg1*Δ⍺, for which *n* = 20. For all other calculated mutation rates, *n* = 10 was used for each strain. Mutation rates and confidence intervals for all fluctuation assays in this study are provided in Table S6. All raw data and mutation frequencies are included in Table S7.

Mutation frequencies were calculated by determining the mean number of colonies on YPD plates, adjusting the YPD mean value to the dilution and volume of cells plated to the selective medium (Adjusted YPD mean = YPD mean × ((volume of cells plated to selective media × dilution factor of cells plated to YPD) / (volume of cells plated to YPD × dilution factor of cells plated to selective media)), and dividing the number of resistant colonies on each plate by the adjusted YPD mean. Unlike mutation rates, mutation frequencies do not take into account plating efficiencies or mutant fitness but enable the display of all data points and their distributions (Figures S13, S14, S15, S16, S17). Figure S13 displays the raw data used to calculated Figure 1B mutation rates. Figure S14 shows the raw data corresponding to the mutation rates in Figure S1. Raw data in Figure S15 corresponds to the mutation rates in Figure S2. Figure S16A displays raw data used to calculate mutation rates shown in Figure 2A; S16B data corresponds to mutations rates in Figure S8. The raw data presented in Figure S17A and S17B were used to calculate mutation rates in Figure 5A and 5B, respectively. Box-and-whisker plots were generated using the ggboxplot command in the ggplot2 package (v3.3.5) in R (v4.1.0) using default parameters.

### Characterizing mutation spectra

Following selection on antifungal drug media, resistant colonies were streak purified to YPD medium. Only one colony from each plate was streak purified for characterization, ensuring each mutation event identified was independent. Genomic DNA was isolated from the purified colonies, and genes in which mutations are known to cause resistance to the corresponding antifungal drug were PCR amplified (*URA5* and *URA3* for 5-FOA-resistant colonies^54, 55^, *FRR1* for rapamycin+FK506-resistant colonies^46, 47^, and *FUR1* and *UXS1* for 5-FC-resistant colonies^48^). Oligonucleotides used for all PCR reactions in this study are listed in Table S8. PCR products were subjected to gel electrophoresis, imaged by the Quantity One® Software, and products of interest were extracted from agarose gels using a QIAgen gel extraction kit and sequenced through classical Sanger sequencing conducted at Genewiz. *FRR1* was PCR amplified in a total of 39 H99, 77 Bt65, and 37 Bt81 R+F^R^ independent colonies; 37 H99, 31 Bt65, and 27 Bt81 *FRR1* PCR products were sequenced. *URA3* and *URA5* were PCR amplified in 5 H99, 11 KN99⍺ *msh2*Δ, 5 Bt65, and 5 Bt81 5-FOA^R^ independent colonies; *URA5* and *URA3* PCR products were sequenced from 5 H99, 2 Bt65, and 5 Bt81 strains. *FUR1* and *UXS1* were PCR amplified in 9 H99, 5 Bt65, and 10 Bt81 5-FC^R^ independent colonies; 2 Bt81 *FUR1* PCR products were sequenced. Sequenced mutations, including transposon insertions, were characterized with both Sequencher software and the Clustal Omega Multiple Sequence Alignment program^80^. Identified transposon insertion sequences in *FRR1*, *URA5*, and *FUR1* are listed in Table S9.

### Illumina sequencing

Single colonies from strains for whole-genome Illumina sequencing were inoculated in 50 mL of liquid YPD medium and grown overnight at 30°C, shaking. Cells were collected and lyophilized as previously described^53^, and high molecular weight DNA was isolated following the CTAB protocol as previously described^81^. Strains were barcoded and sequencing libraries were generated with the Kapa HyperPlus library kit for 300bp inserts, pooled, and sequenced using paired-end, 2 x 150bp reads on an Illumina HiSeq 4000 platform at the Duke University Sequencing and Genomic Technologies Core facility.

### Generation of F_1_ progeny

Bt65**a** x H99α *crg1*Δ and Bt81**a** x H99α *crg1*Δ F_1_ progeny were generated by genetically crossing either Bt65 or Bt81 with H99 *crg1*Δ on Murashige Skoog (MS) medium (Sigma) following Basic Protocol 1 as described in Sun et al. 2019^82^. Basidiospores were randomly isolated through microdissection after three weeks of incubation on MS following Basic Protocol 2 as described in Sun et al. 2019^82^.

### Nanopore sequencing and genome assemblies

The DNA samples for nanopore sequencing were isolated and purified using the CTAB DNA preparation protocol described previously^83^. The size estimation of the obtained DNA was done using PFGE electrophoresis and quality was determined using NanoDrop. Once the high-quality DNA was obtained, sequencing was performed using the MinION device with the MinKNOW interface. During sequencing, Bt65, Bt89, and Bt133 were multiplexed together whereas six of the Bt65**a** x H99α progeny were multiplexed for a second sequencing run. For multiplexing, samples were barcoded using EXP-NBD103/EXP-NBD104 kits and libraries were made using SQK-LSK109 kit as per the manufacturer’s instructions. The libraries generated were sequenced on R9.4.1 flow cell and reads were obtained in .fast5 format. These reads were then converted to fastq format using Guppy_basecaller (v 4.2.2_linux64). The reads were de-multiplexed using qcat (https://github.com/nanoporetech/qcat) or Guppy_barcoder (part of Guppy_basecaller) with barcode trimming option during processing. Bt81 nanopore sequencing was done as a standalone sample using an R9 flow cell (FLO-MN106) and basecalling was performed during the run itself.

The sequences obtained for each sample were then assembled via Canu (v2.0 or v2.1.1) to obtain contig-level genome assemblies. For the assembly, only >2 kb long reads were used for the Bt65**a** x H99α F_1_ progeny and Bt81, whereas >5 kb were used for Bt65, Bt89, and Bt133 genomes. Contigs were then assigned chromosome numbers based on their synteny with the reference genome, H99. The numbering of chromosomes involved in translocations was assigned based on the respective syntenic centromere. Some of the chromosomes were not fully assembled and were broken into multiple contigs (Chr 1, Chr 2 for Bt65, Chr 2, Chr11, Chr14 for Bt89, and Chr 2, Chr 5 for Bt133). For such cases, the respective contigs were joined artificially and then processed by read-mapping to obtain complete collinear chromosomes. Specifically, the contigs were stitched together in orientation as determined based on their synteny. Corrected reads obtained from Canu were then mapped to the respective genomes and duplicated or missing regions from the junction were identified. The chromosome sequence was then corrected accordingly by inserting/correcting/deleting sequences and a full-length chromosome sequence was obtained. Complete resolution of junctions was obtained for Bt65, Bt89, and Bt133 genomes by this approach. However, some of the Bt65 F_1_ progeny chromosomes could not be resolved, probably due to hybrid origin of sequencing reads, and were left with gaps as such.

Once chromosome level genome assemblies were obtained for the Bt65, Bt81, Bt89, and Bt133 genomes, the genome sequences were further processed to improve telomeric and subtelomeric regions. For this purpose, the corrected reads obtained from Canu were mapped back to the respective chromosome-level genomes using minimap2 v2.14. The obtained bam files were then analyzed manually by IGV and consensus or, in a few cases, individual reads (up to 30 kb) representing extra sequence beyond an assembled chromosome were extracted as sam files. These consensus extra sequences were then added onto the chromosome sequences to obtain longer chromosomes. In some cases, read mapping also resulted in the identification of incorrect sequence assembly at subtelomeric regions, and in those cases, the sequence was trimmed until a consensus sequence was observed at the end of the chromosome. Once these corrections were made, the genome assemblies were polished via one round of nanopolish and five rounds of pilon, except for the Bt81 genome, for which only 5X pilon polishing was performed. As a result of these corrections and polishing, final assemblies were obtained for each of the four isolates and are described in the study. For the Bt65 F_1_ progeny genome assemblies, the subtelomeric extension/curation was not performed, but they were polished using both nanopolish and 5X pilon.

### Centromere, telomere, and Cnl1 mapping

Centromeres in Bt65, Bt81, Bt89, and Bt133 were defined based on their synteny with the reference H99 genome (genome assembly ASM301198v1) The final polished genomes were used and centromere locations were identified by BLASTn analysis using H99 centromere-flanking genes as query sequences. Once the centromere locations were defined, Tcn1-Tcn6 locations within those regions were mapped by BLASTn analysis. For the representation, only BLAST hits longer than 400 bp were mapped. For the overlapping BLAST hits with multiple Tcn elements, the longest and best BLAST result was used, and the rest of the matches were discarded from further analysis. All the hits were then visualized using Geneious Prime and maps were exported as .svg files, which were then processed using Adobe Illustrator.

For the Cnl1 mapping at the subtelomeres, the longest *CNL1* insertion sequence from the Bt65 genome was used as the query sequence and BLASTn was performed against each genome. BLAST hits longer than 50 bp were mapped to the respective genomes and visualized using Geneious Prime where the hits were color-coded based on their lengths. The zoomed views for these maps were then exported as .svg files, processed using Adobe Illustrator, and combined with centromere Tcn mapping analysis to generate final figures.

RepeatMasker was used to annotate all transposons in the *de novo* genome assembly of Bt65. For this purpose, RepeatMasker (v4.0.7) with Dfam (v3.3) and RepBaseRepeatMaskerEdition-20181026 libraries was used, supplemented with RepBase EMBL database (v26.04)^84–86^. The “-species fungi” option was used to identify all repeats in the genome and provided additional support for the manual Tcn and Cnl1 mapping.

### Synteny maps

Synteny comparisons between the genomes were performed using SyMAP v4.2 with the H99 genome as the reference (genome assembly ASM301198v1). The synteny comparison was conducted using default parameters and synteny block maps were exported as .svg files. The maps were processed using Adobe Illustrator for visualization. The phylogenetic relationship as depicted in Figure S6 was drawn based on the earlier representation^20^. The telomere and centromere locations were marked manually based on the presence of the telomere repeat sequence and Tcn mapping, respectively.

For the centromere comparisons, all centromere sequences along with Tcn annotations were converted into GenBank format. The files were then used for synteny comparison via EasyFig v2.2.3. The maps were exported as .svg files which were processed in Adobe Illustrator.

### Recombination maps for Bt65 x H99 F_1_ progeny

Six of the Bt65**a** x H99α F_1_ progeny were sequenced with on the nanopore MinION sequencing platform and their genomes were assembled and polished using the methods described above. Once their genomes were assembled, recombination maps were generated by mapping the Illumina sequence data from the parental strains to each of the progeny genomes. For this purpose, both H99 and Bt65 Illumina reads were used from published datasets (SRR642222 and SRR647805 for H99; SRR836876, SRR836877, SRR836878, SRR836880, SRR836884, and SRR836885 for Bt65). Reads from all runs were merged to obtain a single file for both H99 and Bt65. The reads were then mapped to the progeny genomes using Geneious Prime default mapper with three iterations. Variants with 90X coverage and at least 90% variant frequency were called from these mapped files. These variants along with coverage analysis were then used to identify recombination sites and generate recombination maps. Cnl1 mapping for each of progeny genome was performed as described above. The location of *ZNF3* in each genome was identified by BLASTn analysis using H99 *ZNF3* (CNAG_02700) as the query sequence.

### Genetic variant calling and segregant filtering

Whole-genome sequencing data of 28 F_1_ progeny from the Bt65**a** x H99α *crg1*Δ cross were aligned via BWA (v0.7.12-r1039)^87^ to an H99 reference genome (downloaded from FungiDB [http://fungidb.org/fungidb/] on April 15^th^, 2020; FungiDB-46_CneoformansH99_Genome.fasta) and genetic variants between Bt65 and H99 were called using SAMtools (v0.1.19-96b5f22941)^88^ and FreeBayes (v1.2.0)^89^. Approximately 300,000 raw genetic variants were identified across the segregants. The genotypic correlation between progeny, the read coverage per genetic variant, and the ratio of reads suggesting the H99 vs. Bt65 allele per variant were monitored across the genome to identify clones, progeny with aneuploid genomes, and heterozygotic diploids (respectively). Two pairs of clones were identified (Supplementary Table S10) and one segregant from each pair was retained for analysis. F_1_ progeny 25 was identified as a heterozygotic diploid (Supplementary Figure S18) and removed from initial analysis. Instances of aneuploidy (and partial duplications) are observed along Chromosomes 3, 4, 11, and 13 within six segregants from this cross and for initial filtering and analysis, those with heterozygotic aneuploidy were removed from analysis (Supplementary Table S10).

### Genetic variant filtering

After removing clones and samples with aneuploidy or diploidy, raw genetic variants were filtered by limiting sites to bi-allelic SNPs, called across all the progeny (100% call rate), with greater than 10X read coverage (and a maximum of 200X), a minor allele frequency of 5%, and a quality score greater than 4 (and less than 5.4). These filtering criteria were selected after examining the bivariate relationships between allele frequency, read depth, and quality scores per chromosome (Supplementary Figure S19A and 14B). Genetic variant sites were also removed if within one kb of the centromere along a given chromosome^33^. After filtering, a total of 215,411 bi-allelic SNPs were retained for further analysis. The median distance between contiguous SNP sites is 45 bp, and less than 0.01% of neighboring sites had a distance larger than two kb. The allele frequencies across the genome ranged between 25 and 75% of segregant with the Bt65 allele, except for a large portion of Chromosome 13, between 0 to 500 kb, where over 88% of segregants inherited the Bt65 allele (Figure S19C). With these data, a Poisson regression (methods described in Roth *et al*. 2018^90^) was used to relate the average number of crossovers across F_1_ progeny as a function of chromosome size. Briefly, for the Poisson regression, the following model was used: log(E(# of crossover⎹ x)) = -b + m × x, where x is chromosome size.

In the Bt65 x H99 F_1_ progeny, this model was estimated as: log(E(# of crossover⎹ x)) = −0.3955 + 0.6415 × x. This model rejected the null hypotheses of a zero intercept (b) and chromosome coefficient (m), with *p*-values of 0.0162 and 2.68 × 10^-9^, respectively. The model predicts an obligatory 0.673 crossovers per chromosome, which increases by a ratio of 1.899 per Mb increase in chromosome size. With this model, the estimated genome-wide, physical-to-genetic distance in this cross is 8.14 kb/cM.

### QTL mapping

For use in association tests, across 24 F_1_ progeny and the two parental strains, the 215,411 bi-allelic SNPs were collapsed into 1,237 unique haplogroups made up of genetic variants sites that co-segregated within the segregant genomes, such that, between any two haplogroups, at least one segregant contains a change in allele (i.e. a recombination event between the Bt65**a** and H99α *crg1*Δ genomes). Collapsing genetic variants into haplogroups reduces the number of repeated tests in association mapping and computational costs^91^. Across the 1,237 haplogroups, the mutation rate on rapamycin + FK506 medium was used as the phenotype. The phenotypic data was assumed to be non-normal based on visual inspection, and a Shapiro-Wilk test for normality (two-sided) confirms this assumption (Shapiro-Wilk-statistic = 0.822, *p*-value < 0.000527). Because of this, we used a two-sided Kruskal-Wallis H-test for genome wide QTL mapping.. The –log_10_ (*p*-value) from this test across haplogroups was monitored to identify QTL. Significance thresholds were established via 10,000 permutations with an α = 0.01 as described in Churchill and Doerge (1994)^92^, and 95% confidence intervals for the QTL locations were generated as described in Visscher *et al.* (1996)^93^. No adjustments for multiple comparisons are needed in QTL mapping as a permutation based threshold was used to establish genome-wide significance The heritability at the peak of identified QTL was estimated using linear regression (with 1 degree of freedom) with the model: *M* = μ + β*I* + *e*, where *M* is the mutation rate x 10^7^, *e* is an error term, μ is the mean mutation rate x 10^7^, *I* is an indicator variable for the allele at the QTL peak – coded as 0 if from H99α *crg1*Δ or 1 if from Bt65**a** – and β is the effect of having the H99α *crg1*Δ vs. the Bt65**a** allele at the QTL. The variation explained (R^2^) from this model was used as an estimate of the heritability. The calculated R^2^ value had an associated F-statistic of 43.57 and a *p*-value or 7.94 × 10^-7^.

### Gene annotation and SNP effect prediction

For genes within the identified QTL spanning Chromosomes 3 and 11, the alleles between H99 and Bt65 were imputed using filtered SNP data (described above). The published H99 reference strain annotation (downloaded from FungiDB [http://fungidb.org/fungidb/] on April 15^th^, 2020; FungiDB-46_CneoformansH99.gff) was used to predict changes in protein sequence between the H99 and Bt65 parental backgrounds.

### CRISPR-mediated genetic editing

To change the single nucleotide responsible for the nonsense mutation in the first exon of *ZNF3*, a thymine (base 976004 of H99 Chromosome 3 (CNA3 assembly, accession GCA_000149245.3)), to the wild-type cytosine found in H99 and other phylogenetically closely related strains, as well as to genetically delete *AGO1* and *RDP1*, the Transient CRISPR-Cas9 Coupled with Electroporation (TRACE) system was used^58^. Briefly, the gene encoding Cas9 was PCR amplified from plasmid pXL1-CAS9-HYG. For the *ZNF3* replacement strains, *SH1*-*NEO* construct encoding *NEO* (G418 resistance) targeted to a safe haven locus (SH1) was amplified from plasmid pSDMA57^94^. The *SH1-NEO* construct was linearized with the AscI restriction enzyme (NEB). A 2,171bp region was also amplified from Bt133, containing the wild-type C nucleotide in *ZNF3* exon 1 and no other mutations relative to Bt65 for integration at the *ZNF3* endogenous locus in Bt65 (1,197bp upstream of the *ZNF3* start codon to 971bp after the start codon). To genetically delete *AGO1* and *RDP1*, ∼1 kb regions upstream and downstream of the ORFs of both genes were PCR amplified from Bt65, and the *NAT* dominant marker conferring nourseothricin resistance was amplified from the pAI3 plasmid^95^. Overlap PCR was then used to generate two deletion constructs with the 5’ and 3’ 1-kb regions of homology flanking *NAT* for *AGO1* and *RDP1*. For the sgRNA expression construct, the U6 promoter was amplified from XL280α gDNA, and the sgRNA scaffold was amplified from plasmid pYF515^96^. Overlap PCR was used to generate the sgRNA construct with the U6 promoter and sequences targeting either SH1, the nonsense mutation in *ZNF3*, *AGO1*, or *RDP1*. 2µg of the Bt133 *ZNF3* recombination template, 2µg of the *SH1*-*NEO* linearized construct, 250ng of the *ZNF3* gRNA, 250ng of the SH1 gRNA, and 1.5µg of Cas9 DNA were transformed simultaneously into Bt65 via electroporation using a BIO-RAD Gene Pulser.

*ZNF3*-replacement transformants were selected on YPD + G418 agar plates. Successful transformants were identified through restriction enzyme digest with BtsI-v2 (NEB), which cleaves the first exon of Bt65 *znf3* at the nonsense mutation but does not cleave the first exon of *ZNF3* in Bt133 (or H99) (Figure S20A). PCR was also used to ensure that no transformants had integrated copies of the gene encoding Cas9 or the gRNA constructs and that only a single Bt133 *ZNF3* allele had integrated correctly at the endogenous *znf3* locus (Figure S20B-F). Sanger sequencing was used to further confirm correct replacement of the Bt65 *znf3* allele including the nonsense mutation with the Bt133 *ZNF3* allele. No identified Bt65+*ZNF3* transformants also had a stably integrated copy of the *NEO* gene at the safe haven locus. Transformants lacking *AGO1* or *RDP1* were selected on YPD + nourseothricin agar medium, and successful genetic deletion was confirmed by PCR (Figure 21A). PCR also confirmed integration of the genetic deletion constructs at the endogenous loci as well as integration of only a single deletion construct (Figure 21B-F).

### sRNA isolation and sequencing

*C. neoformans* cells were grown overnight in 50 mL YPD medium at standard laboratory conditions. Following culture, cells were spun down, supernatant was removed, and cells were frozen at −80°C for one hour. Cells were then freeze dried with a Labconco Freezone 4.5 lyophilizer overnight. 70 mg of lyophilized material was used for sRNA isolation following the mirVana miRNA Isolation Kit manufacturer’s instructions. sRNA was quantified with a Qubit 3 Fluorometer and quality was verified with an Agilent Bioanalyzer using an Agilent Small RNA Kit. One biological replicate of each strain was submitted for sRNA sequencing. sRNA libraries were prepared with a QiaPrep miRNA Library Prep Kit and 1 x 75 bp reads were sequenced on the Illumina NextSeq 500 System at the Duke University Sequencing and Genomic Technologies Core facility.

### sRNA data processing

Initial quality control of the small RNA libraries was performed with FastQC 0.11.9^97^ followed by the removal of QIAseq library adapters (5’: GTTCAGAGTTCTACAGTCCGACGATC; 3’: AACTGTAGGCACCATCAAT) with cutadapt 2.8^98^. All untrimmed reads or reads smaller than 14 nt were discarded. The surviving trimmed reads were mapped with bowtie v1.2.3^99^ to the *C. neoformans* Bt65 genome, allowing multiple alignments but no mismatches. The resulting SAM files were converted into BAM file format with SAMtools 1.9^88^ and feature read counts of transposable elements were calculated with BEDTools^100^ using the ‘intersect -w’ option and the annotations of transposable elements, which were identified with RepeatMasker using the repbase database for *C. neoformans*^84, 85^.

Normalization of the read counts to reads per million (RPM) was performed, allowing the comparison of the libraries. Furthermore, the read depth on both DNA strands was analyzed with SAMtools and custom made perl scripts were used to calculate the read size distribution and 5’-nucleotide preference of the small RNA reads as previously described^101, 102^.

### Double-stranded RNA enrichment

For dsRNA enrichment, *C. neoformans* cells were grown overnight in 5 mL liquid YPD medium at 30°C. *Malassezia sympodialis* strains were grown overnight for two days in 5 mL liquid mDixon medium (3.6% malt extract, 1% mycological peptone, 1% desiccated ox bile, 1% Tween 60, 0.4% glycerol) at 30°C. RNA was extracted, and dsRNA was enriched as previously described^69^. dsRNA enrichment in H99, Bt65, and Bt81 did not reveal the presence of any large dsRNA segments (Figure S22). Two biological replicates of each strain were included.

### Biological material availability

All strains and plasmids used and generated in this study are available to others upon request.

## Data availability

All sequencing data is available under BioProject PRJNA749953.

## Code availability

Genetic variant filtering, QTL mapping, and SNP effect prediction was conducted in python (anaconda 3.7.3) via custom scripts available in GitHub (https://github.com/magwenelab/Hypermutator_QTL). All custom Perl scripts reported in the methods for sRNA analysis are also available in GitHub (https://github.com/timdahlmann/smallRNA).

## Acknowledgments

We thank and acknowledge Blake Billmyre for initial project guidance, Shelly Clancey for instruction in conducting fluctuation assays and dsRNA enrichment protocols, Josh Granek for preliminary analyses of hypermutator genomes, Zanetta Chang for assistance in sRNA isolation, Kayla Sylvester for assistance with screening of SDC isolates, and the laboratory of Chris Holley at Duke University for the use of their Nanodrop and BioAnalyzer equipment for preliminary sRNA analysis. We thank Mark Farman and Mostafa Rahnama for stimulating discussion on the impacts of transposons on telomere dynamics. We thank Kevin Zhu for assistance with figure generation. We also thank Sheng Sun, Blake Billmyre, Andy Alspaugh, Sue Jinks-Robertson, Asiya Gusa, and Kayla Sylvester for critical reading and comments on the manuscript. This work was funded by NIH/NIAID F31 Fellowship 1F31AI143136-02A1 awarded to S.J.P. and NIH/NIAID R37 MERIT award AI39115-23, R01 grant AI50113-16, and R01 grant AI33654-04 awarded to P.M.M. and J.H. These studies were supported by a Visiting Professor travel grant awarded by Ruhr University, Bochum, Germany to J.H. J.H. is co-Director and Fellow of the CIFAR program Fungal Kingdom: Threats & Opportunities. We also thank the Madhani Laboratory and NIH grant R01 AI100272 for the KN99α *msh2*Δ deletion strain. T.A.D. and U.K. are funded by the German Research Foundation (DFG) (Bonn Bad-Godesberg, Germany) (KU 517/15-1).

## Author Contributions

S.J.P, V.Y., C.R., T.A.D., U.K., P.M.M., and J.H. designed experiments, interpreted data, and wrote the paper. S.J.P. performed experiments and analyzed fluctuation assay and Sanger sequencing data. V.Y conducted nanopore sequencing and analyzed all resulting data. C.R. and P.M.M. analyzed sequencing data from Bt65 x H99 F_1_ progeny and conducted QTL mapping and analysis. T.A.D. and U.K. analyzed sRNA sequencing data. S.J.P., U.K., P.M.M., and J.H. provided resources.

## Supplementary Figures

**Supplementary Figure S1.**
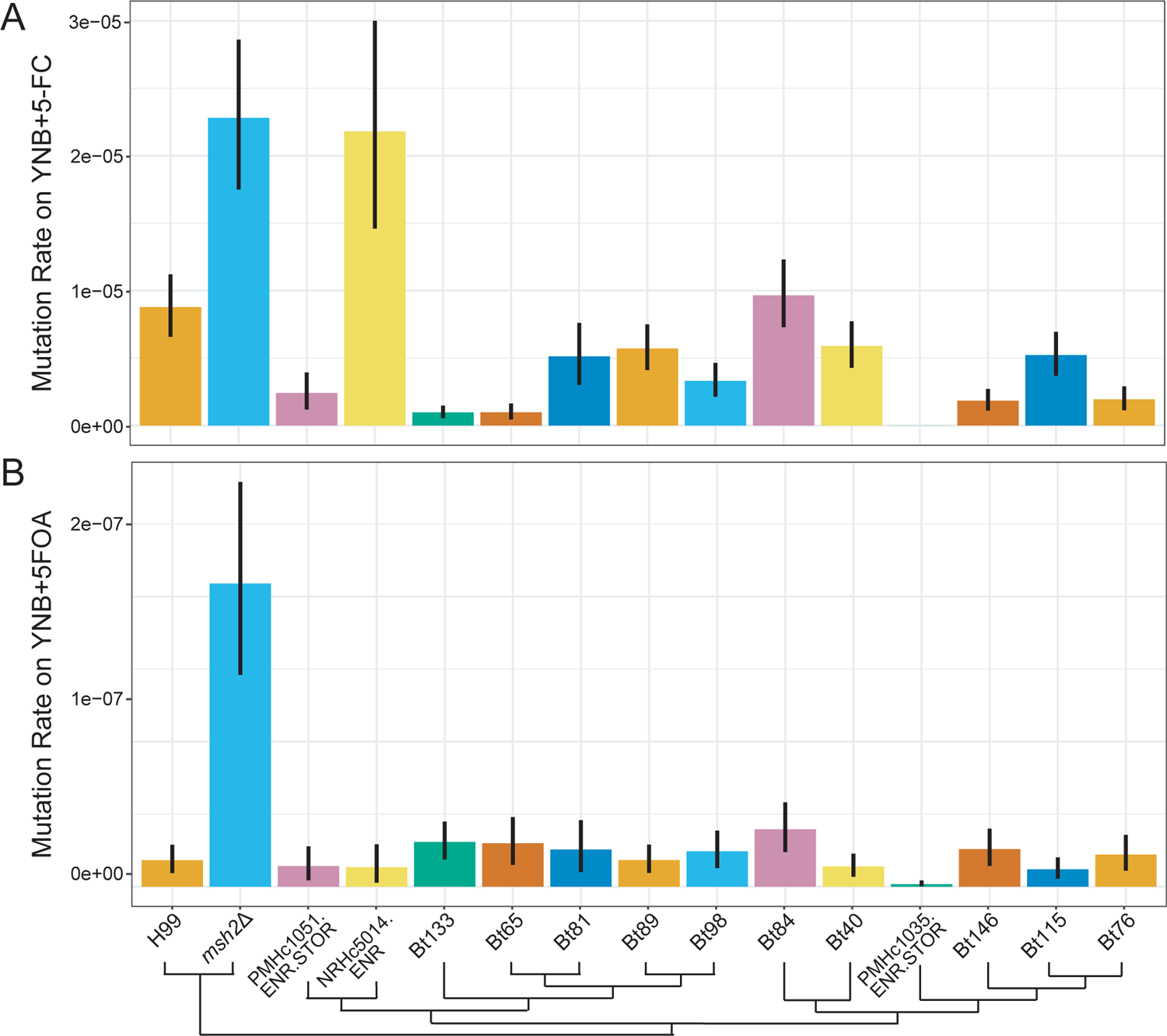
Bt65 and Bt81 do not display a hypermutator phenotype on 5-FC or 5-FOA. Mutation rates of closely related VNBII strains and controls on **(A)** YNB + 5-FC and **(B)** YNB + 5-FOA media. Bars represent the mutation rate and error bars represent 95% confidence intervals; mutation rates represent the number of mutations per cell per generation. Schematic depicts the phylogenetic relationships of all strains included in fluctuation analyses based on Desjardins et al. 2017^20^.

**Supplementary Figure S2.**
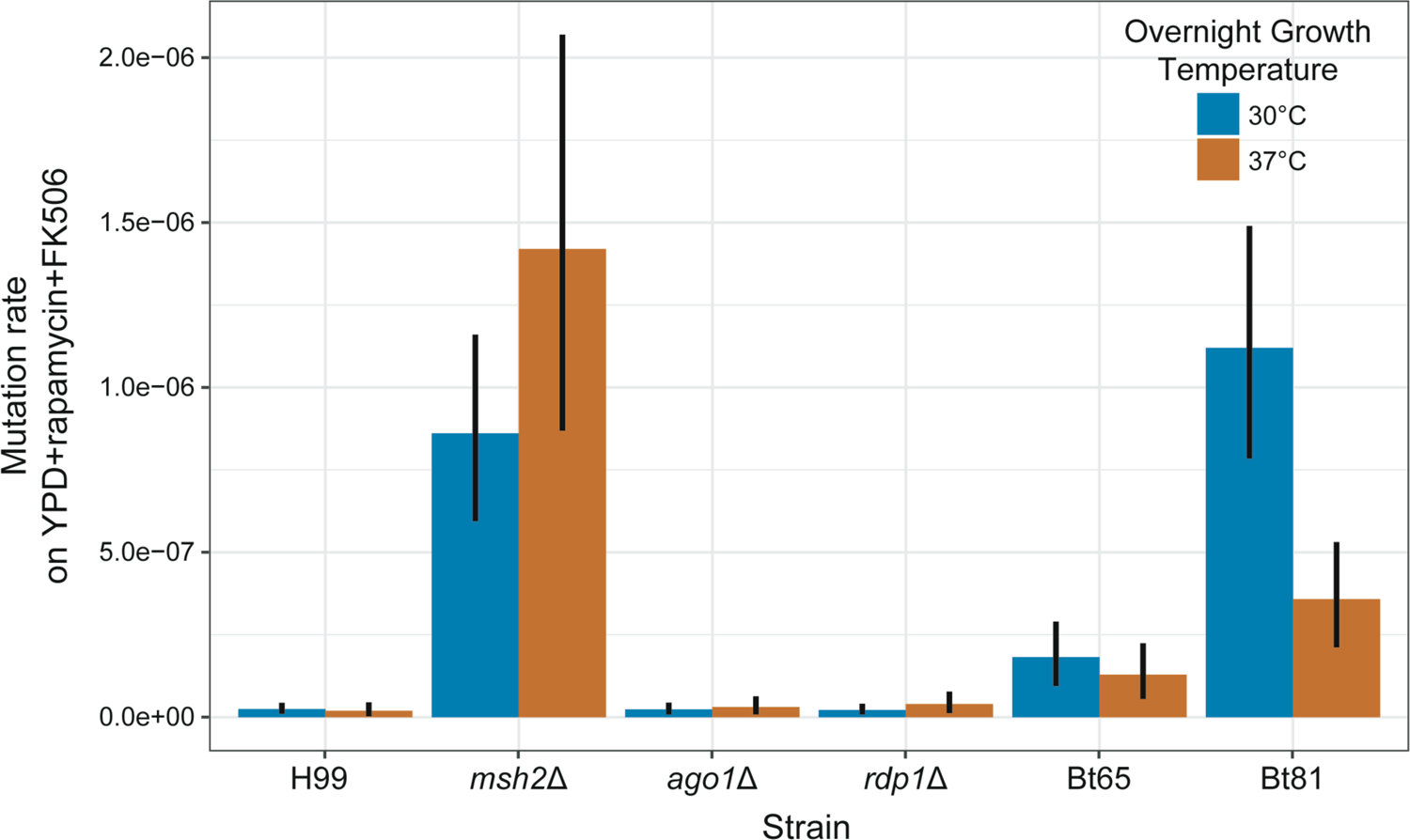
Growth at elevated temperature does not result in increased mutation rates in *C. neoformans* strains. Fluctuation assays were used to quantify the mutation rates of strains grown overnight at 30°C or 37°C and plated on YPD + rapamycin + FK506 medium. Bars indicate mean mutation rate and error bars indicate 95% confidence intervals. Mutation rates represent the number of mutations per cell per generation.

**Supplementary Figure S3.**
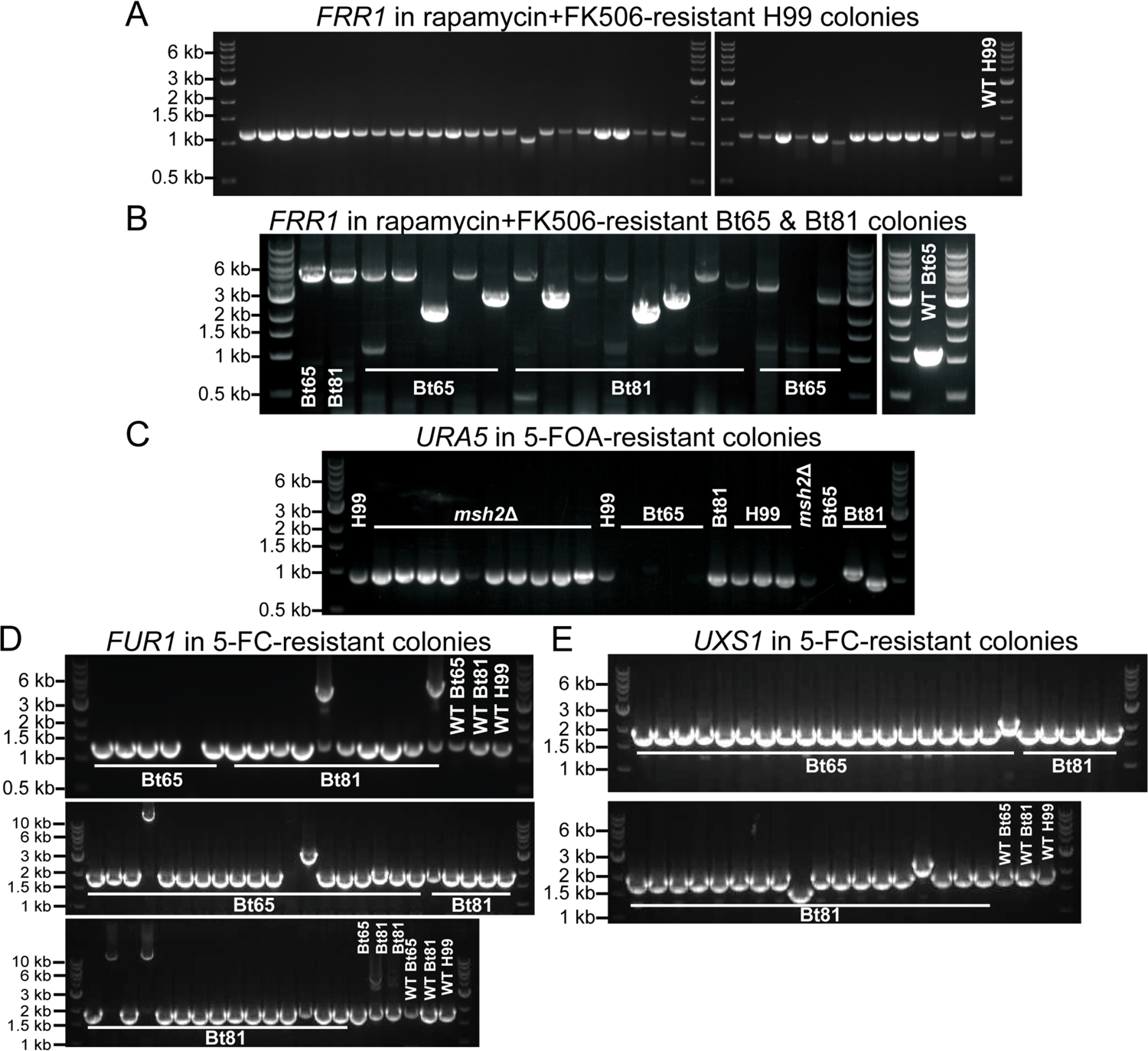
Gel electrophoresis of *FRR1*, *URA5*, and *FUR1* PCR products from resistant colonies. Gel electrophoresis of *FRR1* PCR products from **(A)** all H99 rapamycin + FK506-resistant colonies and a subset of **(B)** Bt65 and Bt81 rapamycin + FK506-resistant colonies sequenced in Figure 1D. PCR amplification of wild-type *FRR1* in *C. neoformans* produces a 1,165 bp electrophoretic species (primers ZC7/8). Gel electrophoresis of a subset of **(C)** *URA5* PCR products from H99, Bt65, and Bt81 5-FOA-resistant colonies and **(D)** *FUR1* and **(E)** *UXS1* PCR products from 5-FC-resistanct colonies of Bt65 and Bt81.

**Supplementary Figure S4.**
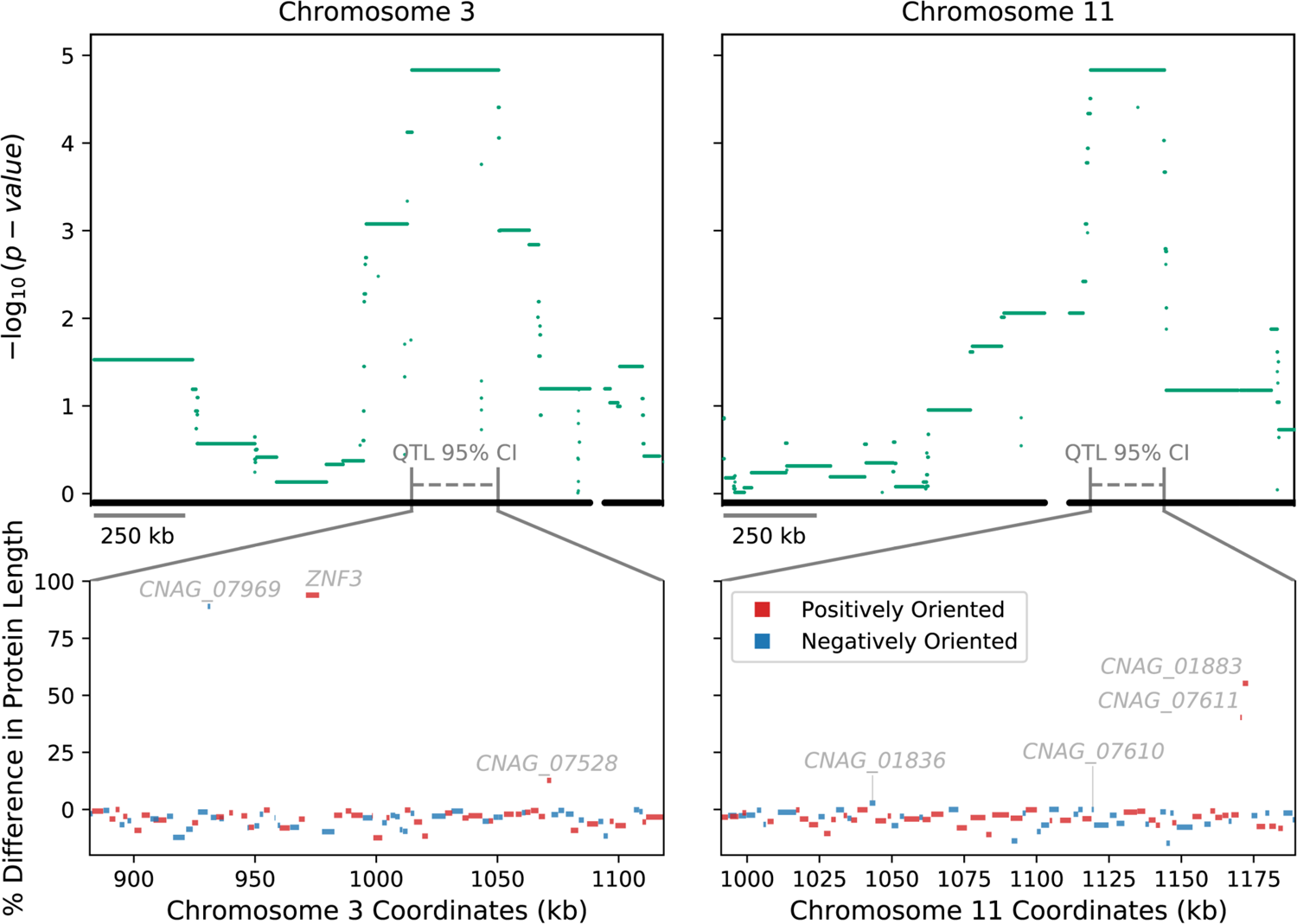
Protein length differences of genes within QTL. In the upper panels, points mark the strength of association (y-axis) between bi-allelic SNP sites and hypermutation for Chromosome 3 and Chromosome 11 (top left and right, respectively). Grey dashed lines depict the 95% confidence intervals (CI) of the two QTL. For the bi-allelic SNPs within the two QTL 95% CIs, *p*-value = 1.46868 × 10^-5^ (Kruskal-Wallis H-test). Lower panels show the predicted differences in lengths of proteins (y-axis) encoded by annotated genes in Bt65 compared to H99 within each 95% CI of the QTL (x-axis) on Chromosome 3 and Chromosome 11 (bottom left and right, respectively). The name of each gene with a predicted nonsense mutation is annotated. Blue and red colors denote the gene orientation.

**Supplementary Figure S5.**
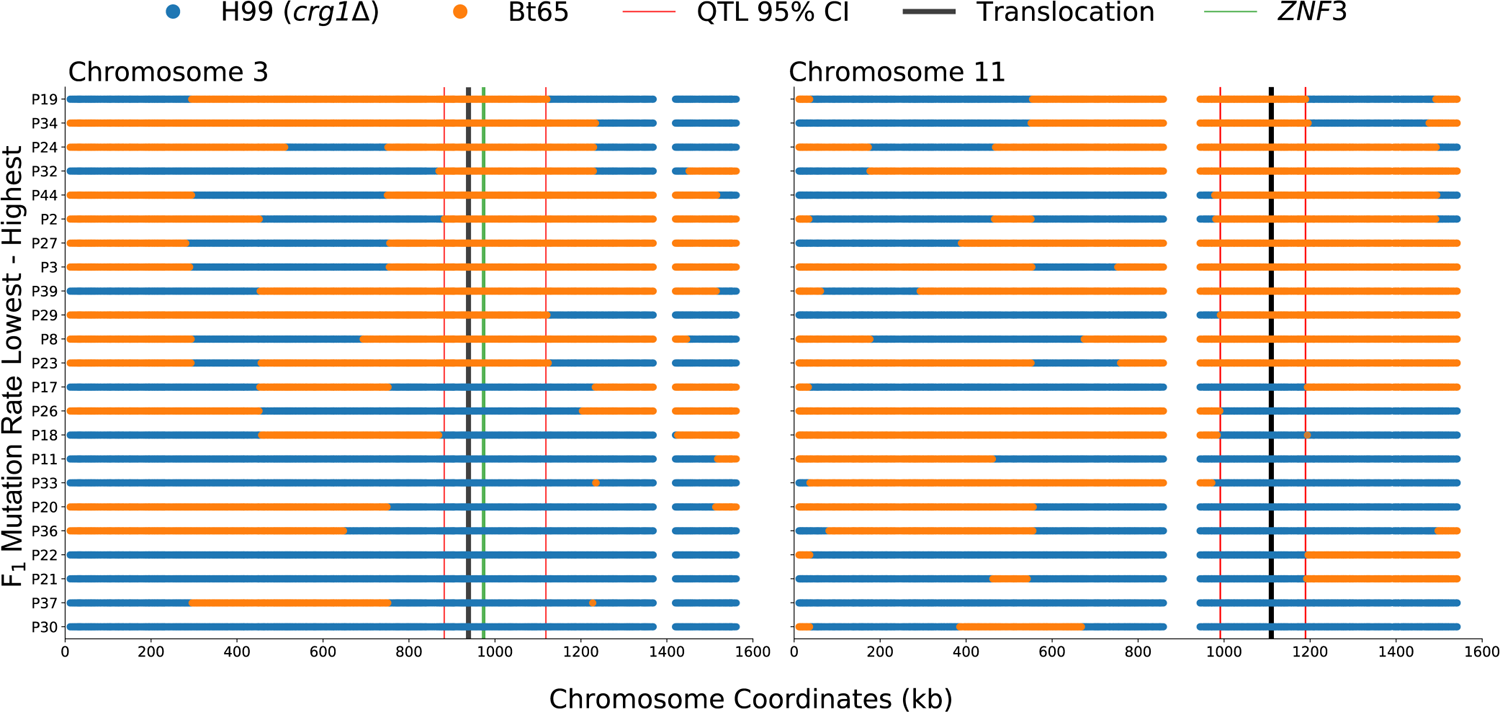
Haplotype maps of Bt65 x H99 F_1_ progeny utilized for QTL mapping. For the QTLs on Chromosome 3 and Chromosome 11 (left and right, respectively) the haplotypes (x-axis) are inferred by SNP data per segregant (y-axis) and colored blue or orange if inherited from H99 *crg1*Δ or Bt65, respectively. Segregants are sorted along the y-axis by the quantification of their mutation rate; largest to smallest, top to bottom. Vertical red lines display the boundaries of the QTL(s). Vertical black lines depict the approximate location of the translocation between H99 and Bt65. The boundaries of the QTG, *ZNF3*, are depicted by vertical green lines. Vertical white spaces indicate the approximate locations of the centromeres.

**Supplementary Figure S6.**
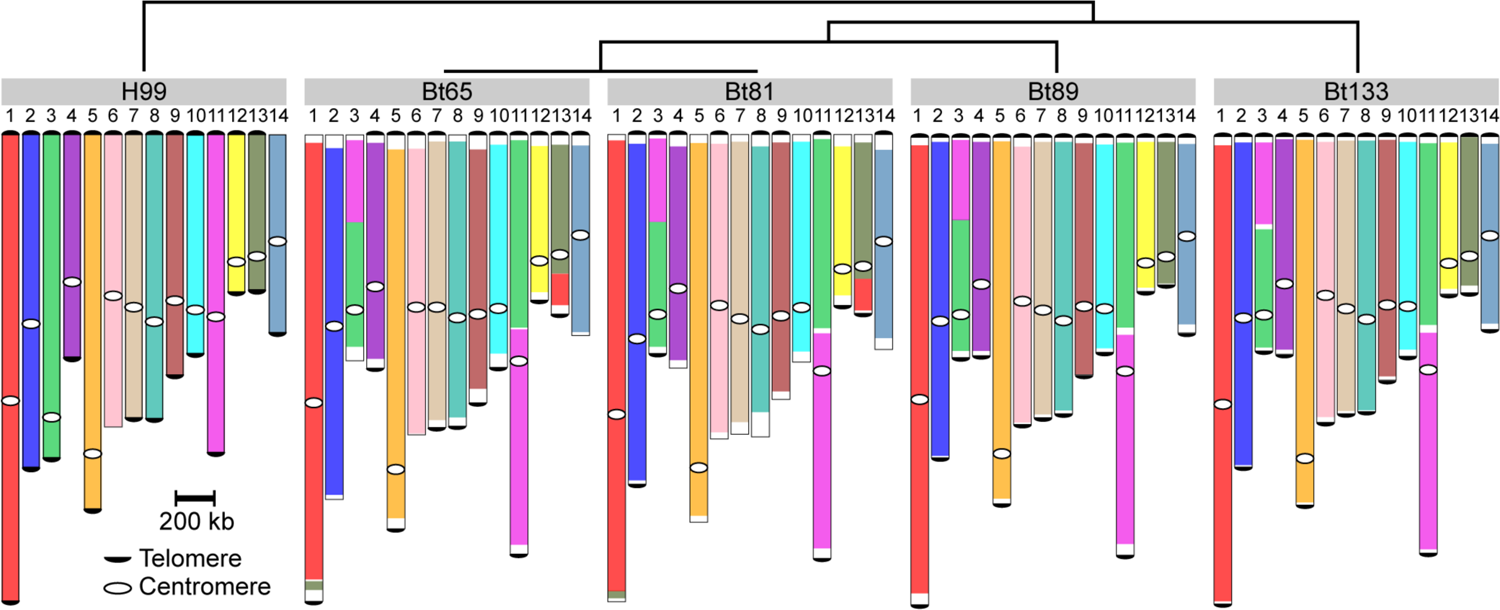
Large-scale genomic rearrangements between the H99, Bt65, Bt81, Bt89, and Bt133 genomes. Nanopore whole-genome sequencing followed by synteny analysis was used to identify all indicated genomic rearrangements with respect to the reference strain H99. There is a chromosomal translocation between Chromosomes 3 and 11 that is unique to H99, and a translocation between H99 Chromosomes 1 and 13 that is unique to Bt65 and Bt81. The phylogenetic relationships of these strains are depicted in the top schematic, telomeric repeat sequences accurately identified in the genomic assemblies are indicated by black half circles, and centromeres are indicated by white circles.

**Supplementary Figure S7.**
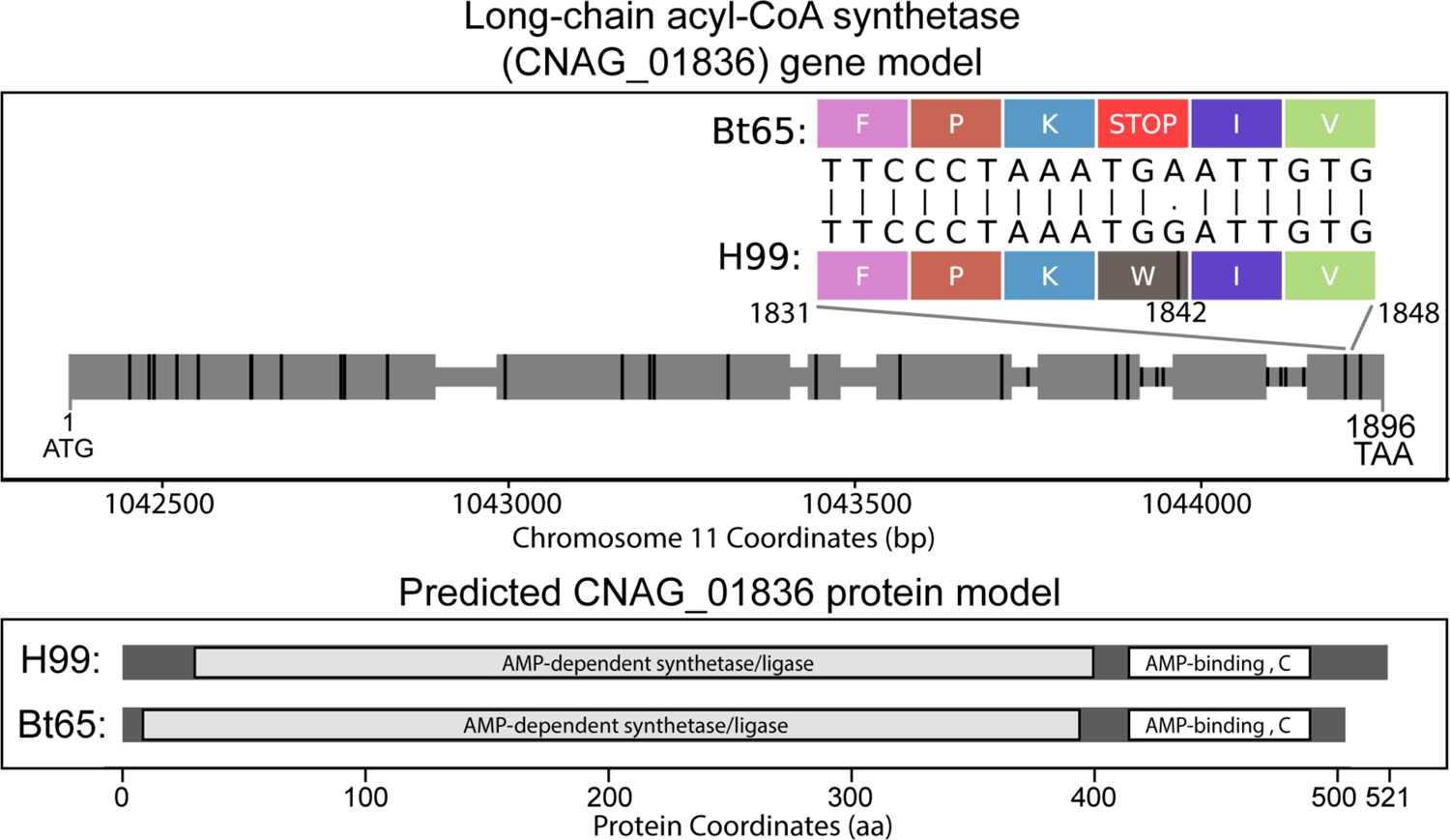
CNAG_01836 gene model. A grey horizontal bar depicts the entire gene body in the upper panel, and larger grey rectangles show locations of exons. The gene is depicted 5’ to 3’, left to right, and is 1896 nt in length. The locations of SNPs differing between Bt65 and H99 are shown by vertical black rungs along the gene model. Amino acids specified by mRNA codons in the indicated region of CNAG_01836 Exon 7 (nucleotide 1831 to 1848) are shown to illustrate the G to A mutation (nucleotide 1842) predicted to cause an early nonsense mutation in Bt65. The bottom panel depicts the predicted outcome of the nonsense mutation on the protein encoded by CNAG_01836 in Bt65.

**Supplementary Figure S8.**
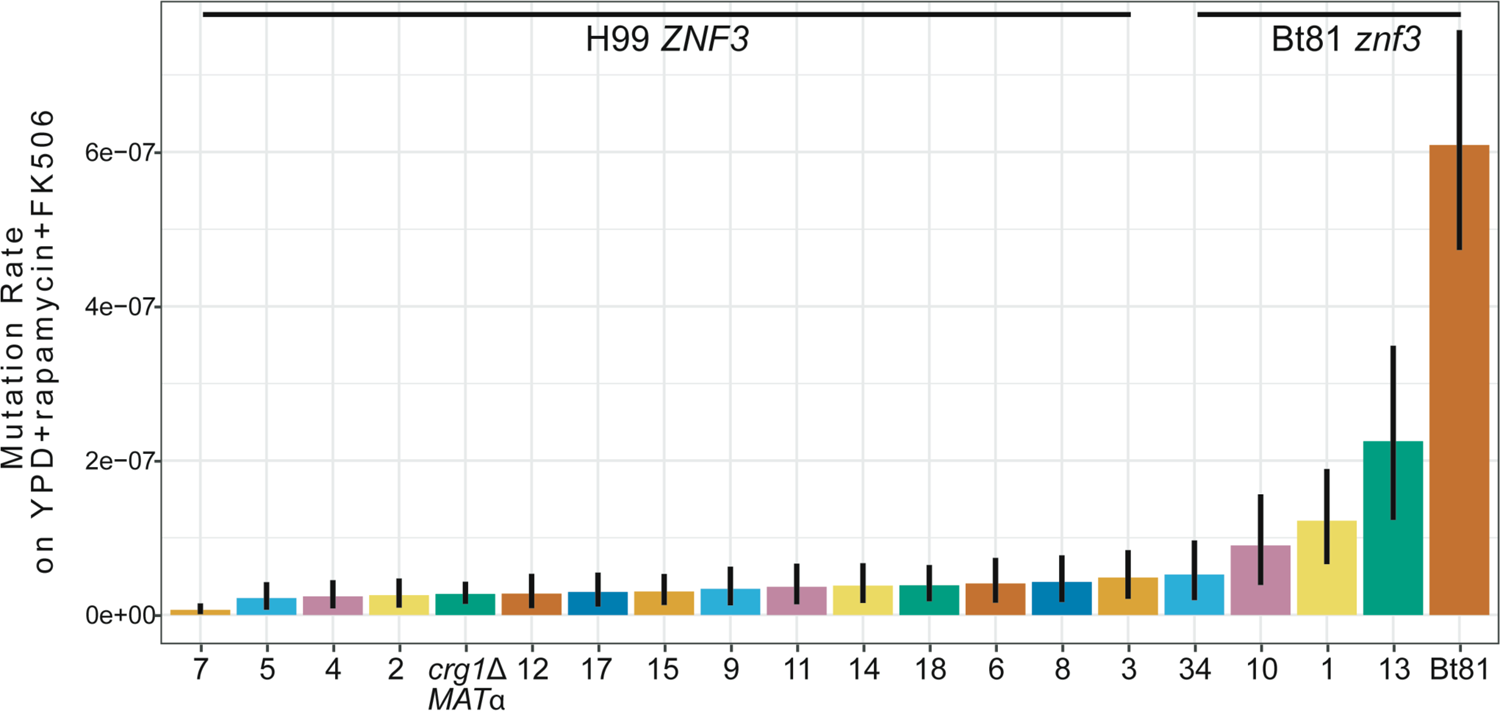
Mutation rates of Bt81 x H99 F_1_ progeny. Fluctuation analysis was used to quantify the mutation rates of the indicated strains on YPD + rapamycin + FK506 medium (y-axis) – sorted smallest to largest, left to right – for F_1_ progeny and the parental strains, H99α *crg1*Δ and Bt81 (x-axis). Bars indicate the mean mutation rate and error bars represent 95% confidence intervals. Mutation rates represent the number of mutations per cell per generation. Inheritance of the Bt81 *znf3* allele or H99 *crg1*Δ *ZNF3* allele in the F_1_ progeny is indicated above mutation rates.

**Supplementary Figure S9.**
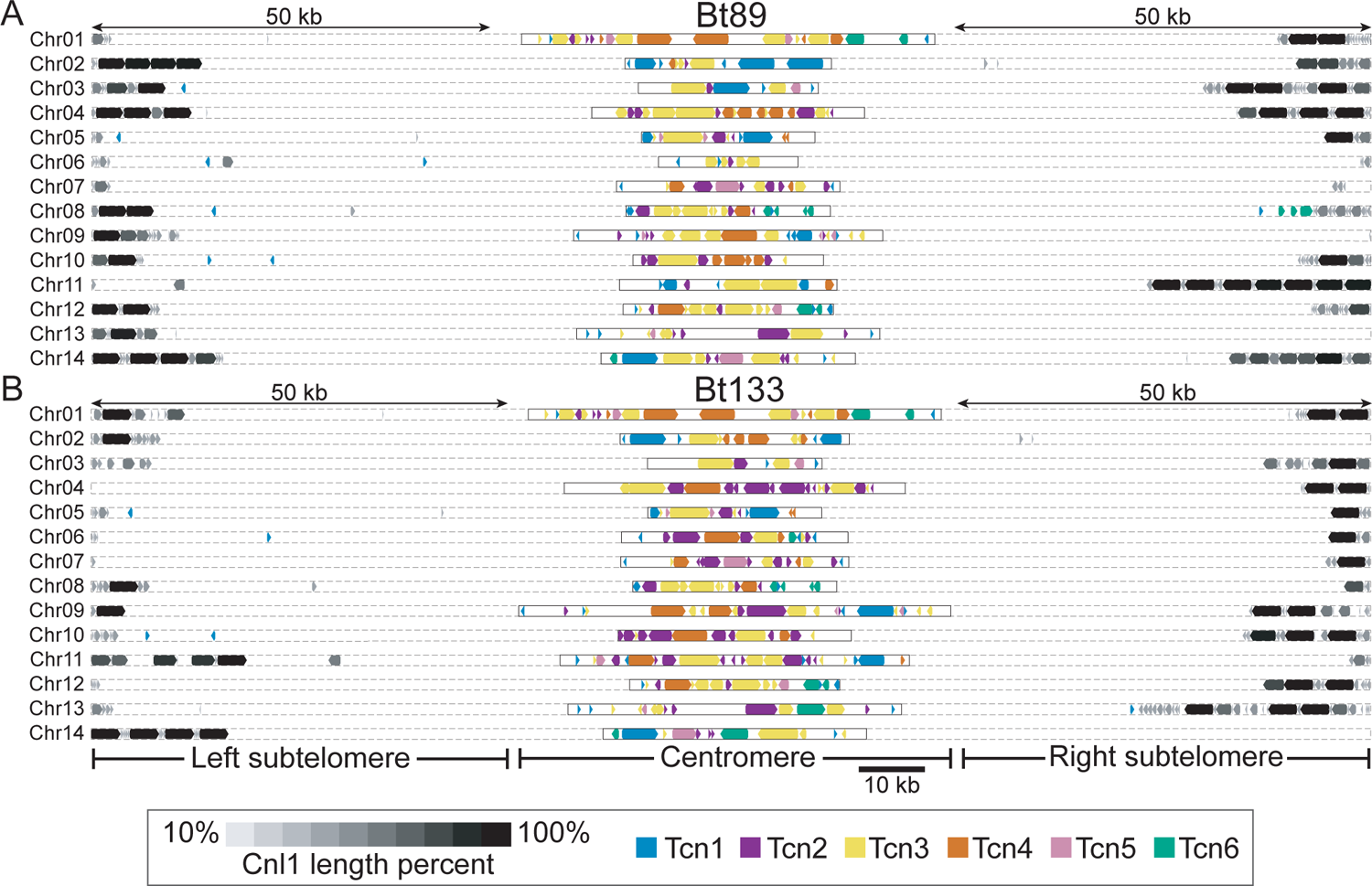
Subtelomeric and centromeric retrotransposons in Bt89 and Bt133. Distributions of the Tcn1-Tcn6 LTR-retrotransposons and the Cnl1 non-LTR retrotransposon in the genomes of **(A)** Bt89 and **(B)** Bt133. 50 kb of subtelomeric regions as well as centromeric regions are displayed for both strains. Shading corresponds to the lengths of the Cnl1 elements, and gene arrowheads indicate the direction of transcription for all retrotransposons.

**Supplementary Figure S10.**
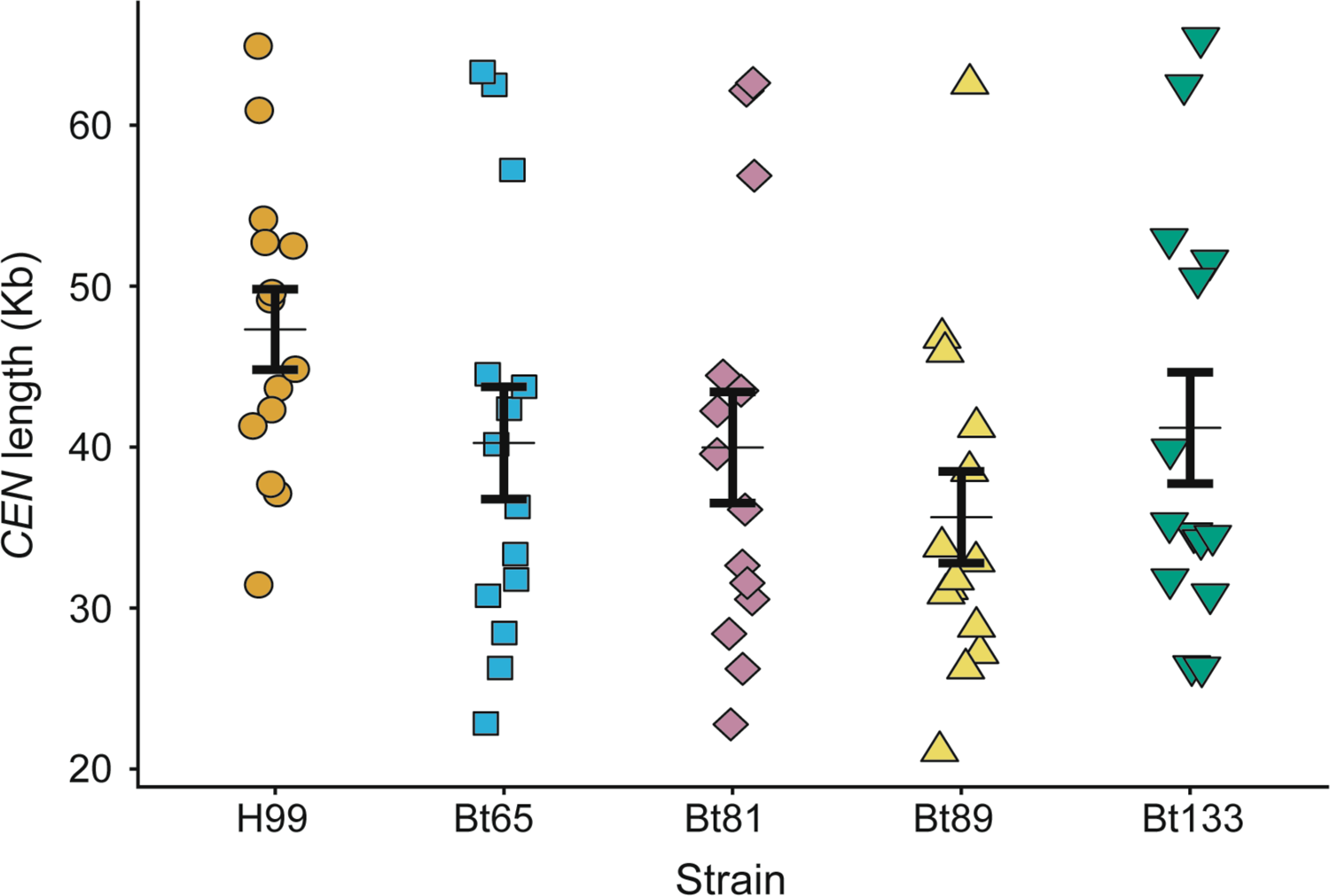
Centromere lengths do not significantly differ among H99, Bt65, Bt81, Bt89, and Bt133. The length of each centromere (y-axis) is plotted for each strain (x-axis). The thin horizontal black line indicates average centromere length and the thicker black error bars indicate the standard error of the mean. No significant difference was found between the average centromere length of each strain (ANOVA, *p*-value = 0.153).

**Supplementary Figure S11.**
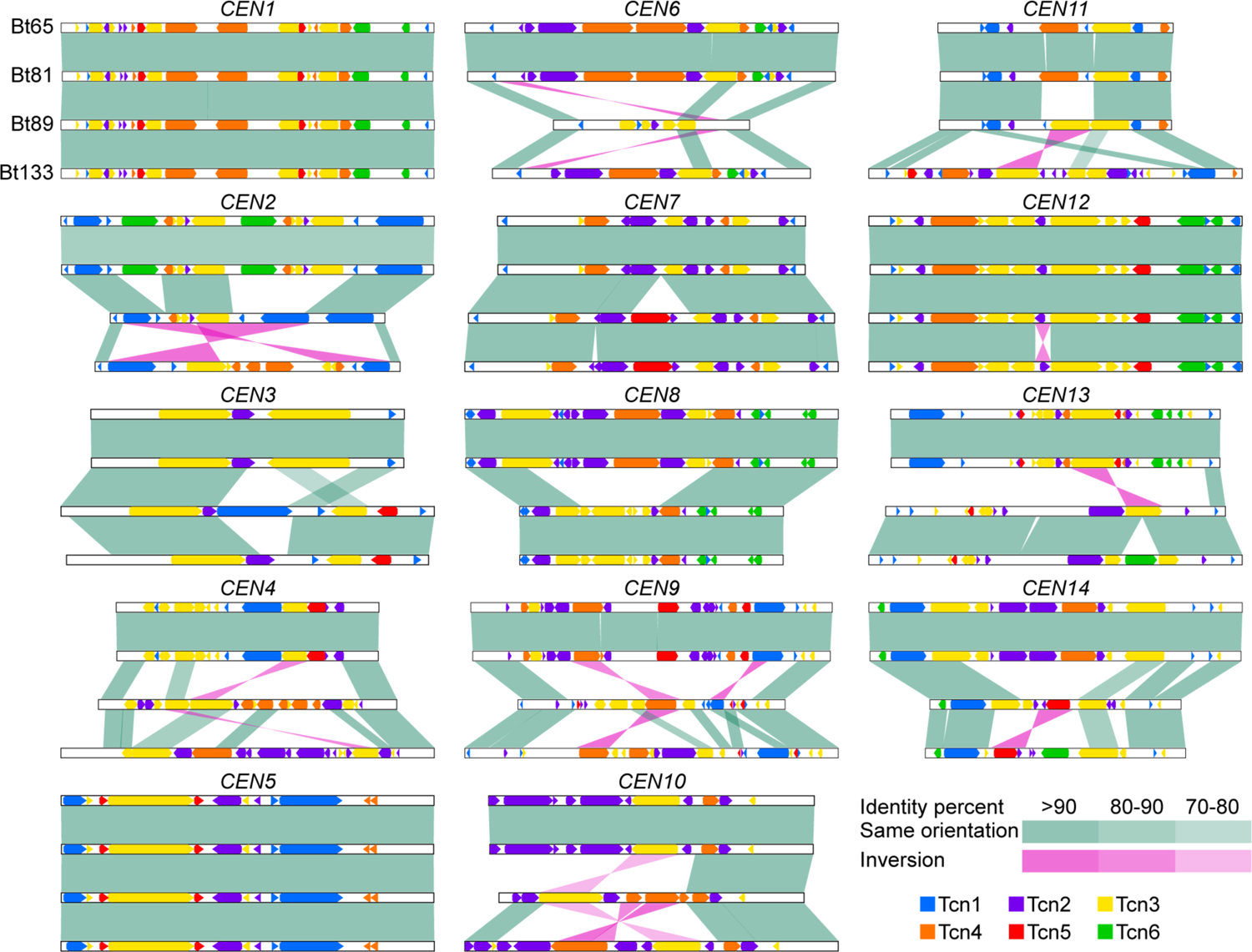
Centromeric rearrangements among hypermutators and two closely related strains. Shared homology, genomic rearrangements, inversions, insertions, and deletions among the centromeres of each of the 14 chromosomes are depicted for Bt65, Bt81, Bt89, and Bt133. The organization of LTR-retrotransposons Tcn1 through Tcn6 are also indicated for each centromere of each strain.

**Supplementary Figure S12.**
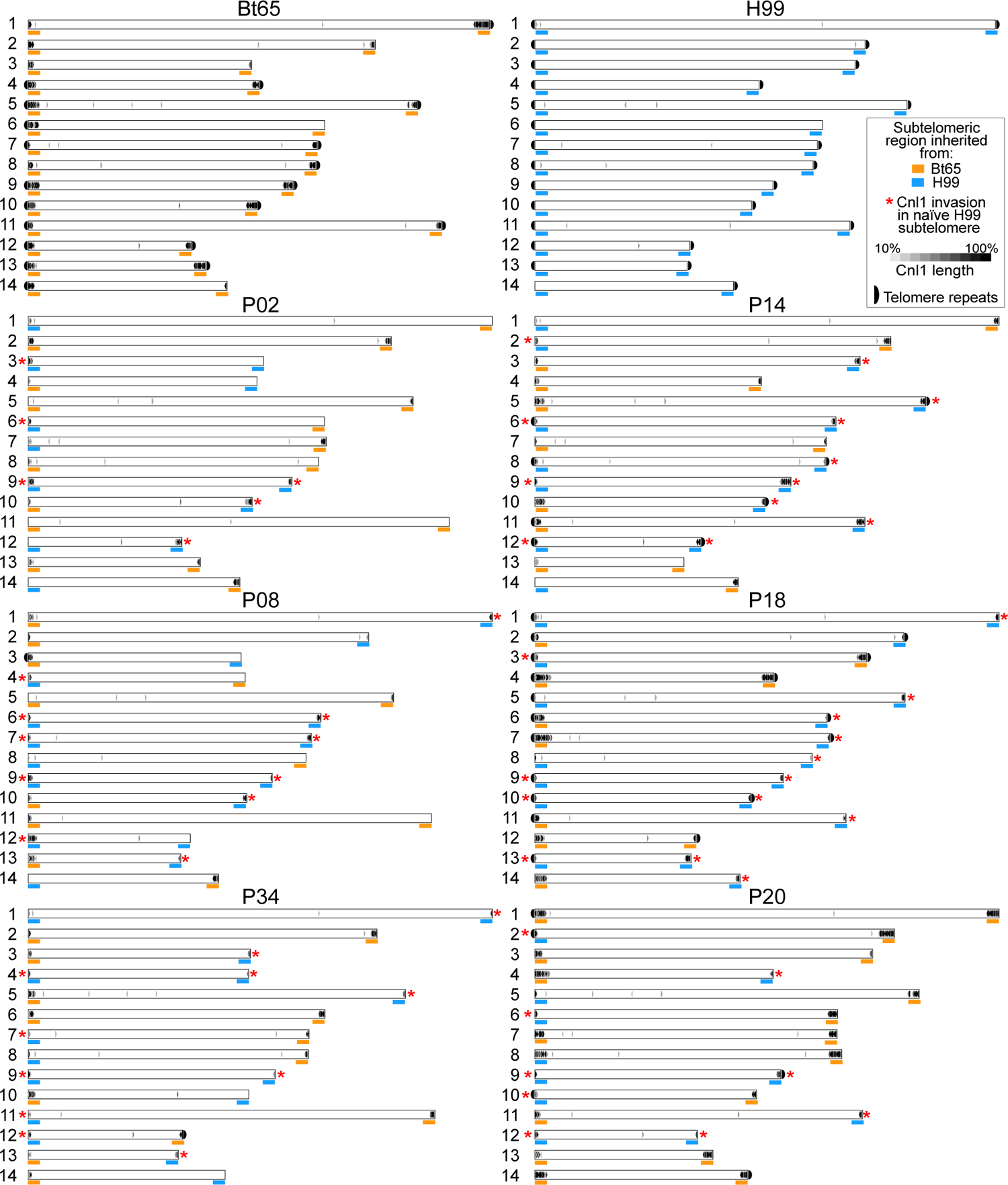
Distribution of Cnl1 among Bt65 x H99 F_1_ progeny and parental strains. The Cnl1 non-LTR elements identified in the nanopore-based whole-genome assemblies are depicted for H99, Bt65, three hypermutator F_1_ progeny (P02, P08, and P34, all on the left), and three non-hypermutator F_1_ progeny (P14, P18, and P20, all on the right). Blue and orange bars under the subtelomeric region of each chromosome indicate which parental strain the region was inherited from (orange for Bt65, blue for H99). Red asterisks indicate invasion of Cnl1 into an H99 subtelomeric region that previously had zero Cnl1 copies/fragments. Accurate assembly of telomeric repeat sequences at the end of each chromosome is indicated by a black half circle. Cnl1 length is also indicated by the shade of black for each element.

**Supplementary Figure S13.**
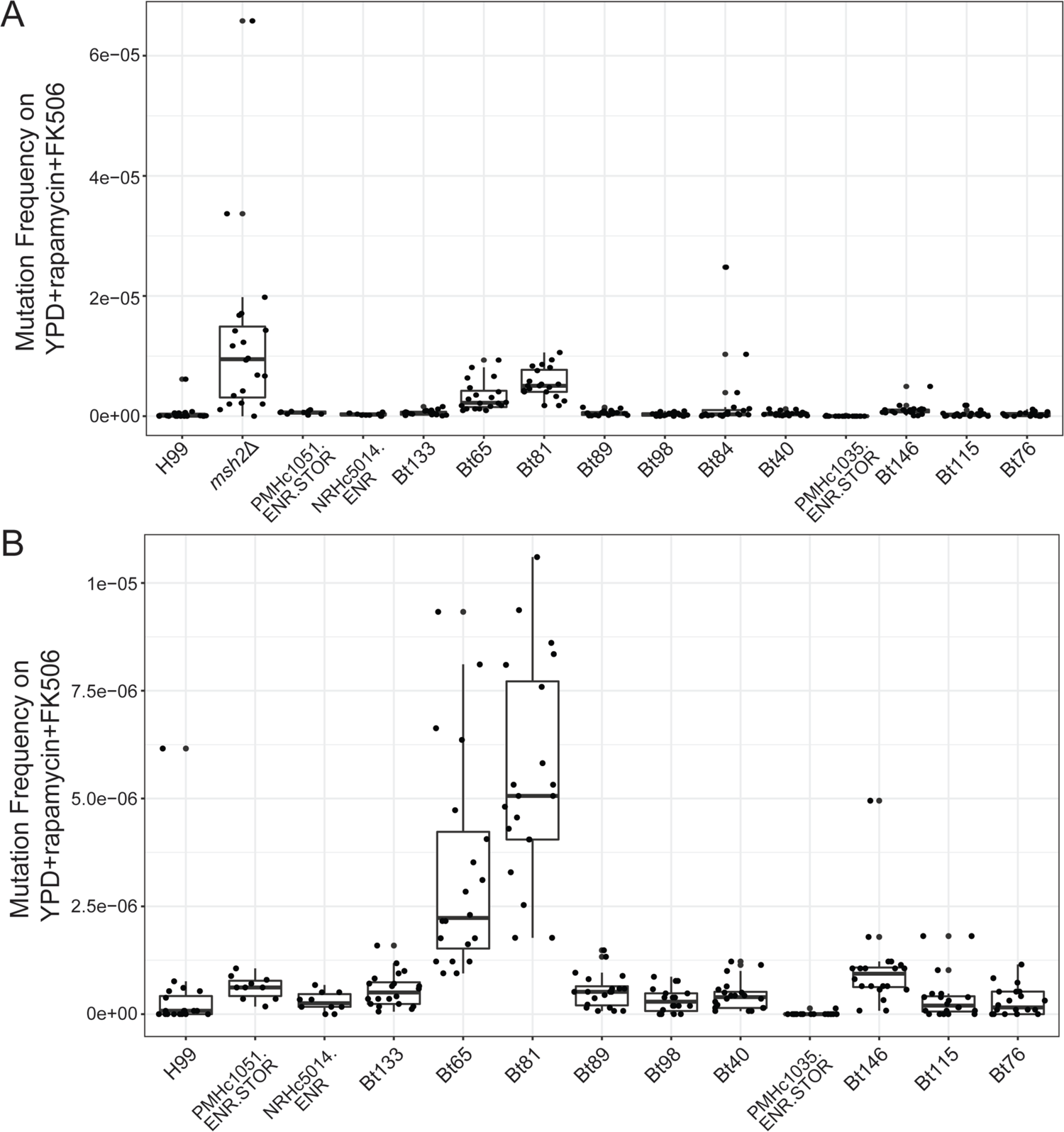
Mutation frequencies of Bt65, Bt81, and phylogenetically closely related strains on YPD + rapamycin + FK506 medium. (A) Mutation frequencies of all strains included in Figure 1B **(B)** Mutation frequencies of all strains excluding the *msh2*Δ mutant positive control and Bt84. In box-and-whisker plots, thicker middle lines in boxes represent the median, the vertical height of boxes represent interquartile ranges (IQRs), whiskers represent 1.5 × IQRs; points above or below the ends of whiskers represent outliers.

**Supplementary Figure S14.**
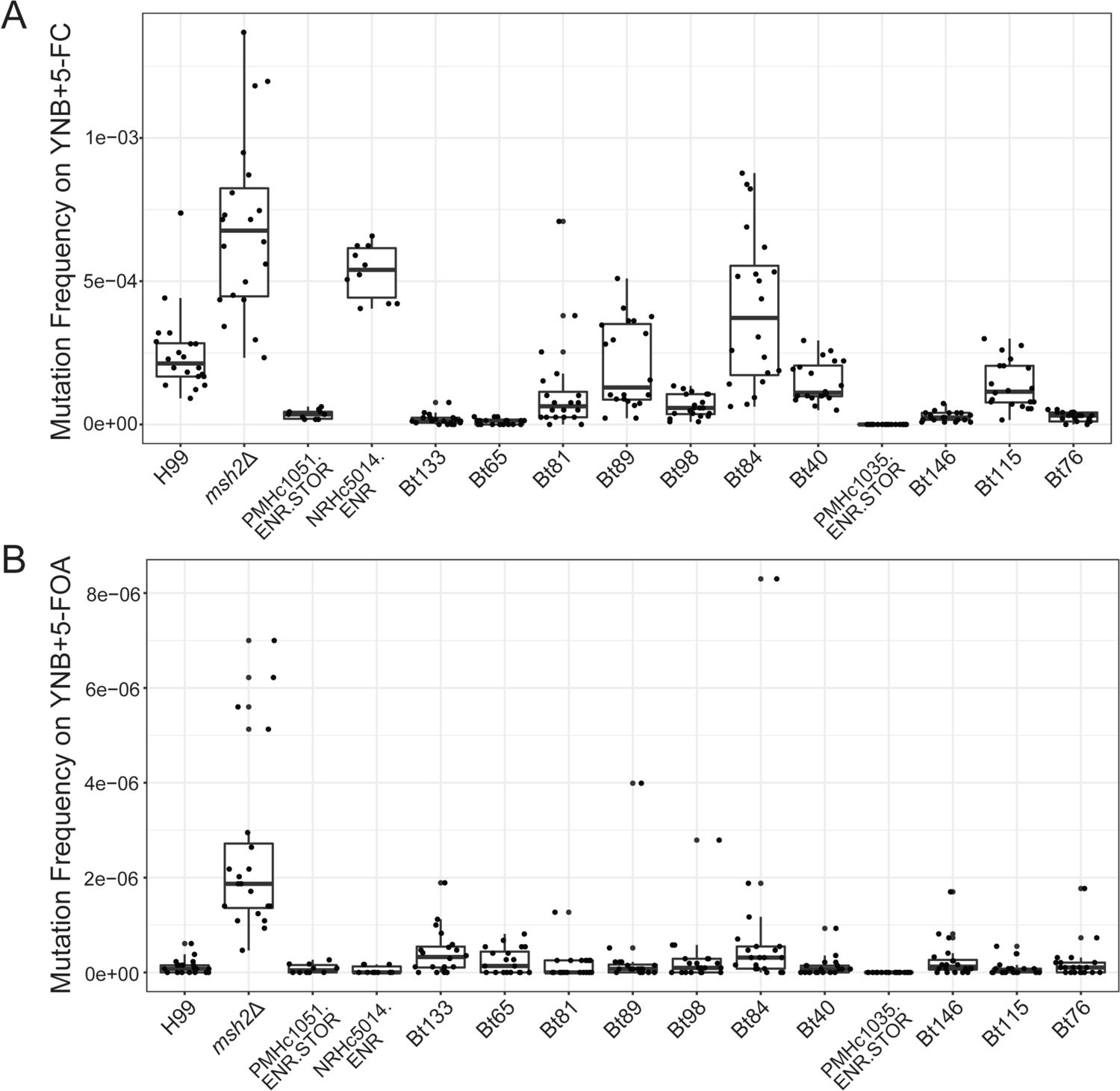
Mutation frequencies of Bt65, Bt81, and phylogenetically closely related strains on 5-FC and 5-FOA. **(A)** Mutation frequencies of all strains included in Figure S1A on YNB + 5-FC medium. **(B)** Mutation frequencies of all strains included in Figure S1B on YNB + 5-FOA medium. In box-and-whisker plots, thicker middle lines in boxes represent the median, the vertical height of boxes represent interquartile ranges (IQRs), whiskers represent 1.5 × IQRs; points above or below the ends of whiskers represent outliers.

**Supplementary Figure S15.**
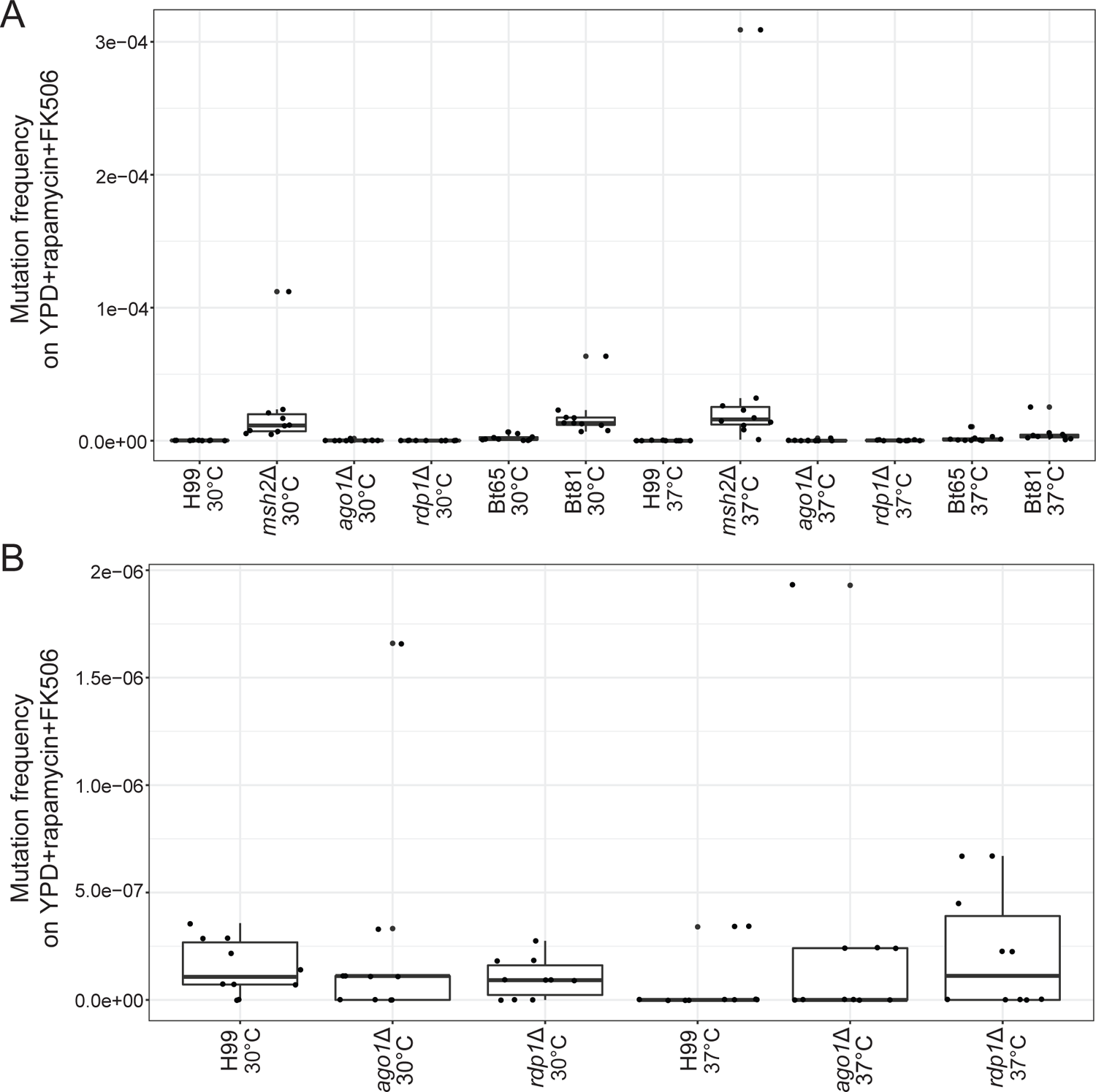
Mutation frequencies of strains grown overnight at 30° or 37° on YPD + rapamycin + FK506 medium. **(A)** Mutation frequencies of all strains included in Figure S2 **(B)** Mutation frequencies of all strains included in Figure S2 excluding *msh2*Δ, Bt65, and Bt81 strains. In box-and-whisker plots, thicker middle lines in boxes represent the median, the vertical height of boxes represent interquartile ranges (IQRs), whiskers represent 1.5 × IQRs; points above or below the ends of whiskers represent outliers.

**Supplementary Figure S16.**
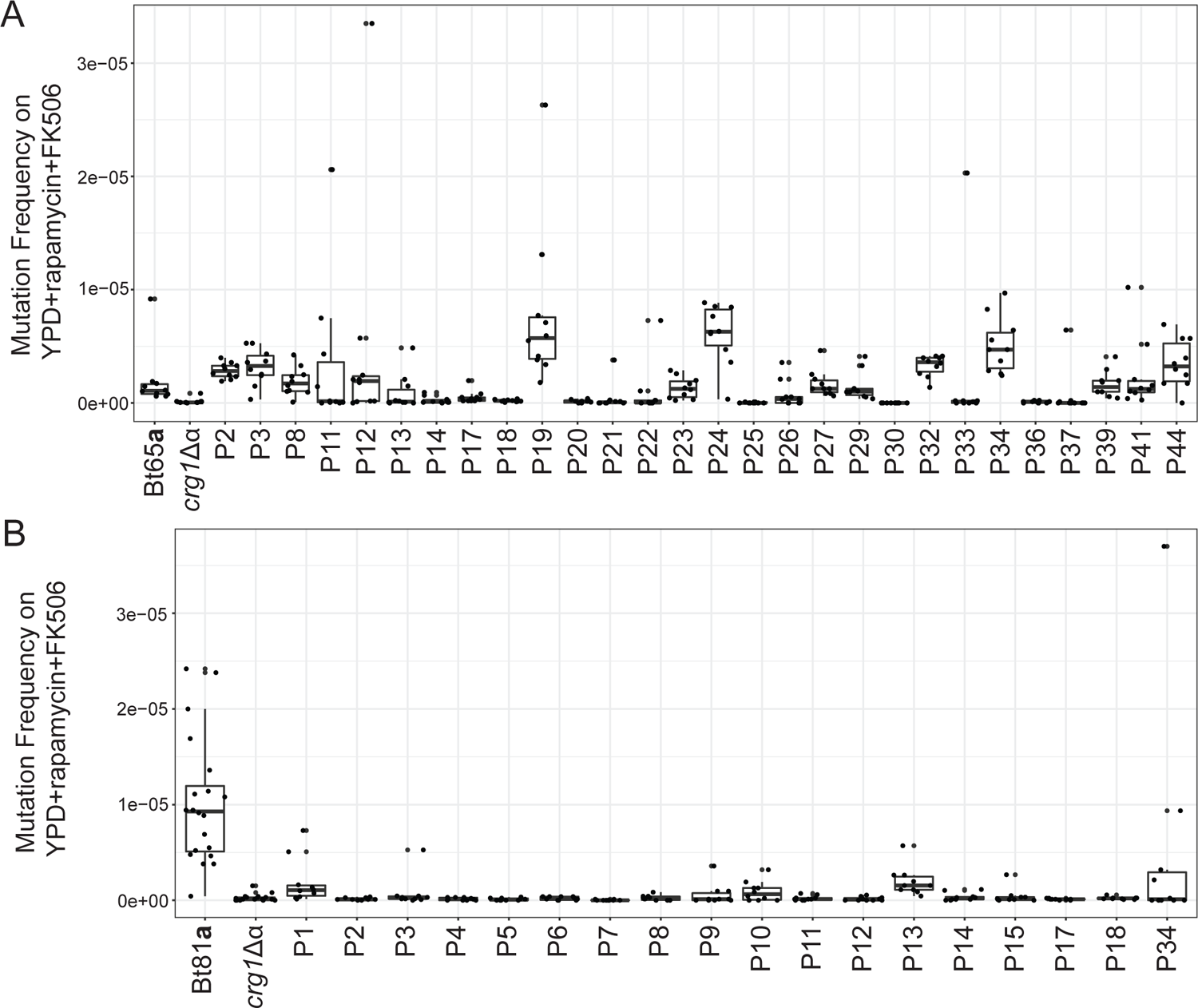
Mutation frequencies of F_1_ progeny derived from hypermutator strains. Mutation frequencies of all **(A)** H99 *crg1*Δ⍺ x Bt65**a** F_1_ progeny included in Figure 2A and **(B)** H99 *crg1*Δ⍺ x Bt81**a** F_1_ progeny included in Figure S8 on YPD + rapamycin + FK506 medium. In box-and-whisker plots, thicker middle lines in boxes represent the median, the vertical height of boxes represent interquartile ranges (IQRs), whiskers represent 1.5 × IQRs; points above or below the ends of whiskers represent outliers.

**Supplementary Figure S17.**
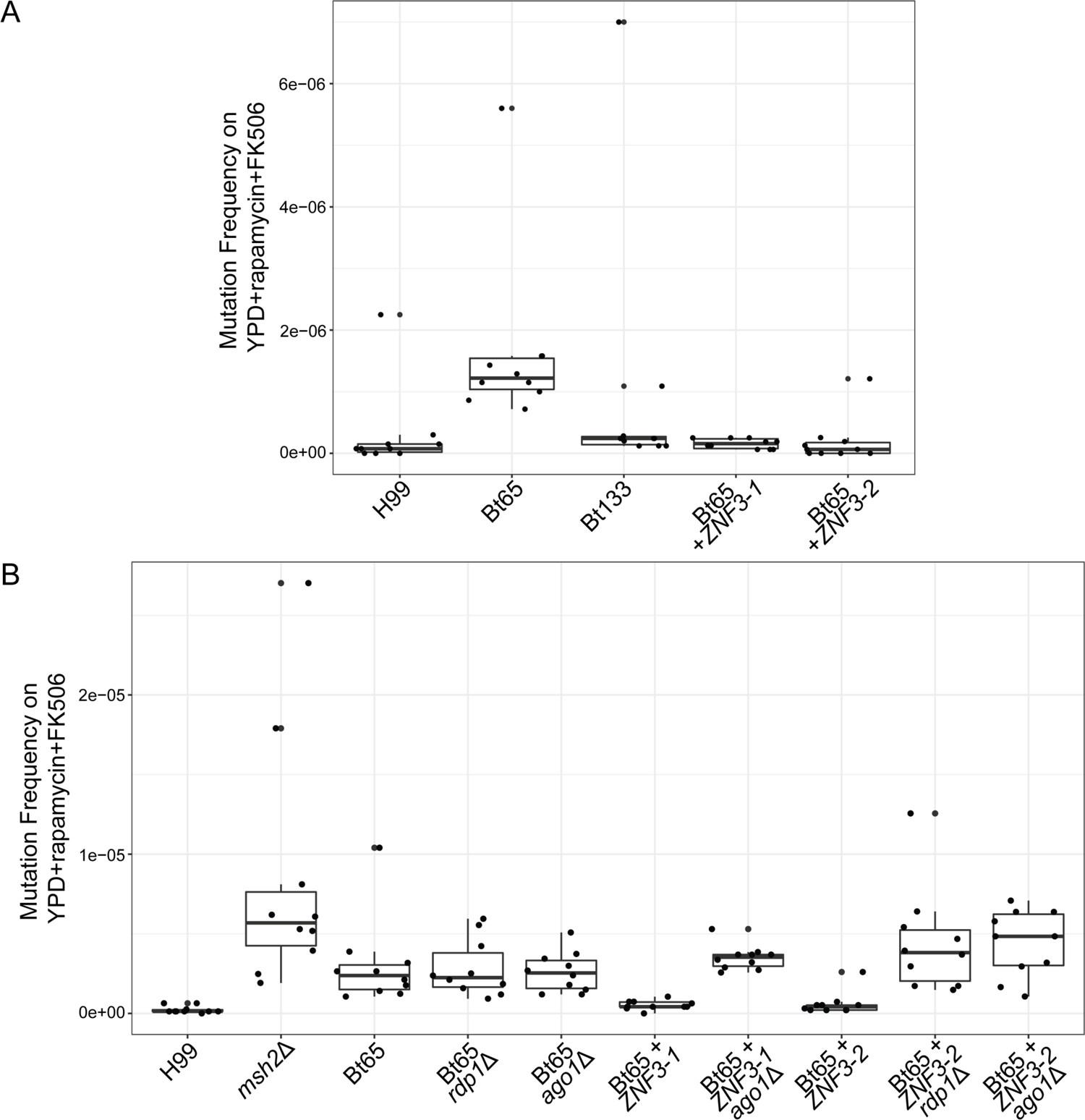
Mutation frequencies of Bt65+*ZNF3*, *ago1*Δ, and *rdp1*Δ genetic mutants on YPD+rapamycin+FK506 medium. Mutation frequencies and raw data used to compute mutation rates shown in Figure 5A **(A)**, and Figure 5B **(B)**. In box-and-whisker plots, thicker middle lines in boxes represent the median, the vertical height of boxes represent interquartile ranges (IQRs), whiskers represent 1.5 × IQRs; points above or below the ends of whiskers represent outliers.

**Supplementary Figure S18.**
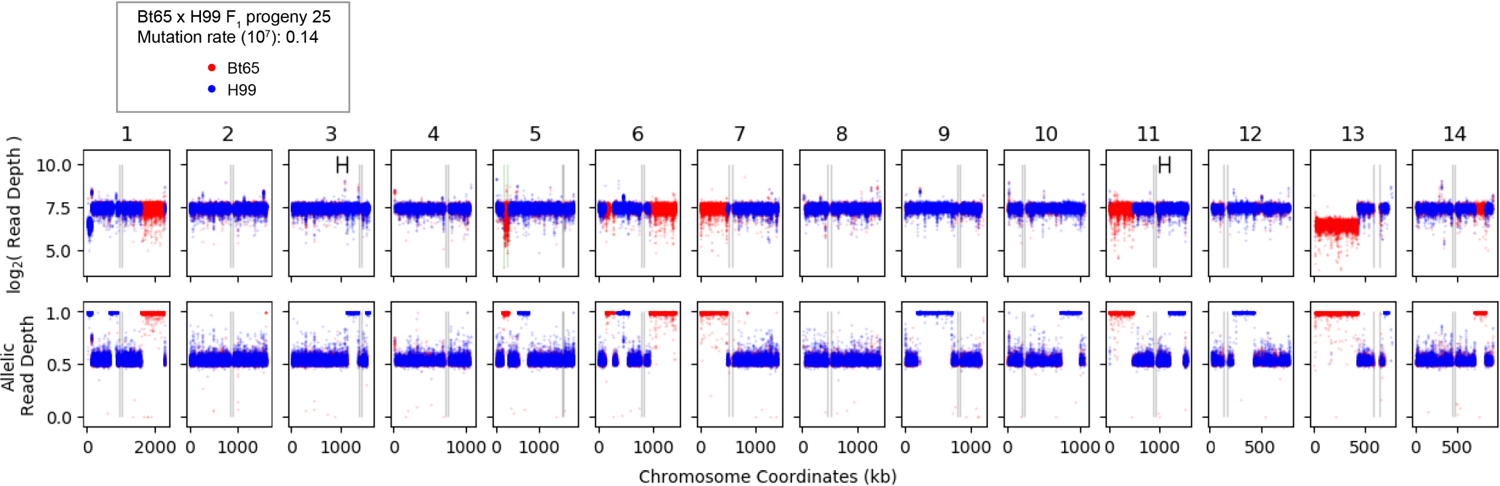
Genomic diagnostic plot of Bt65 x H99 F_1_ progeny 25. For the 14 chromosomes (columns), the log_2_ (read depth) (top row) and allelic read depth ratio (bottom row) per genetic variant are shown for the progeny 25. Red and blue colors indicate the prediction of the allele inherited (Bt65 vs. H99, respectively) at each genetic variant. Allelic read depth ratios nearing 0.5 suggest both alleles are present for a given genetic variant. Marked ranges on Chromosomes 3 and 11 log_2_ (read depth) plots indicate the significant hypermutator QTL. Black vertical lines depict the boundaries of the centromeres. The boundaries of the *MAT* locus on Chromosome 5 are shown by vertical green lines.

**Supplementary Figure S19.**
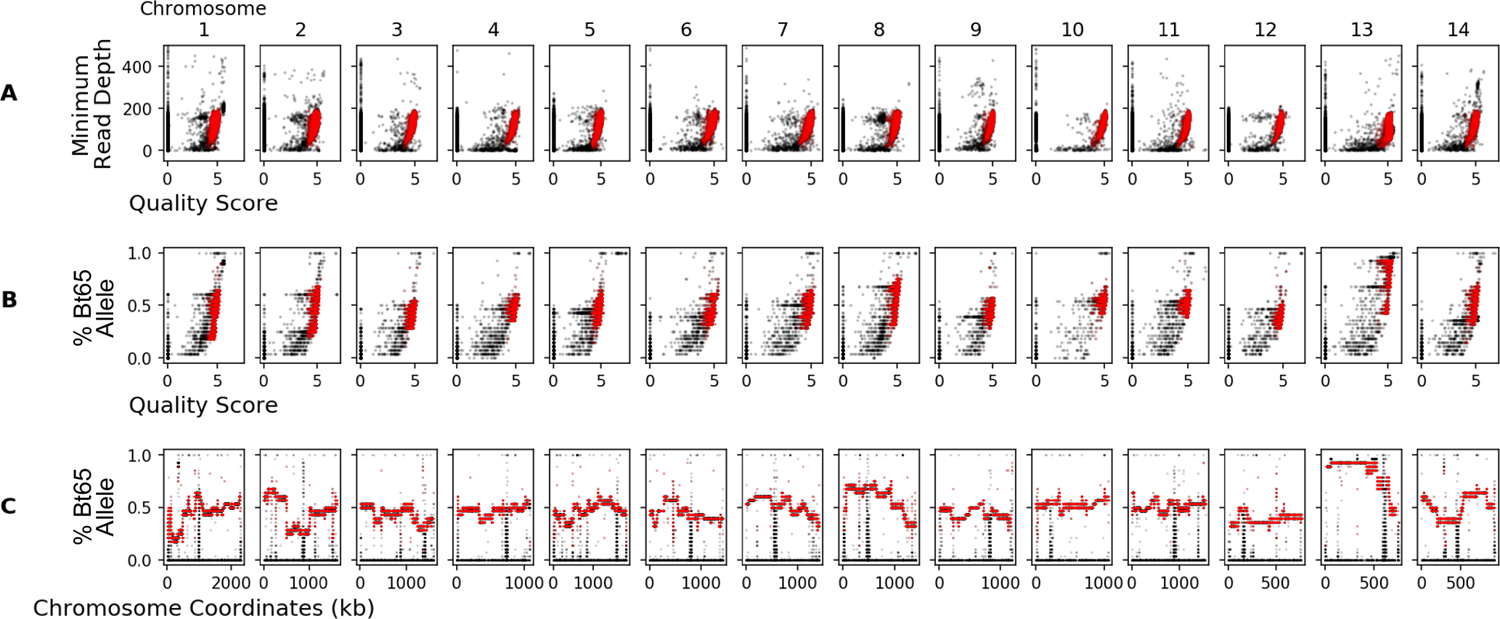
Visualization of genetic variant filtering criteria. (**A**) For the 14 chromosomes (columns), the quality scores of the genetic variants (x-axis) vs. the minimum read depth across the 28 Bt65 x H99 F_1_ segregants (y-axis). (**B**) the quality scores of genetic variants (x-axis) vs. the portion of progeny with the Bt65 allele per genetic variant (y-axis) per chromosome (columns). (**C**) The portion of progeny with the Bt65 allele per genetic variant (y-axis) across each chromosome (x-axis). The raw genetic variants are shown in black and the filtered SNPs, used in analysis, are shown in red.

**Supplementary Figure S20.**
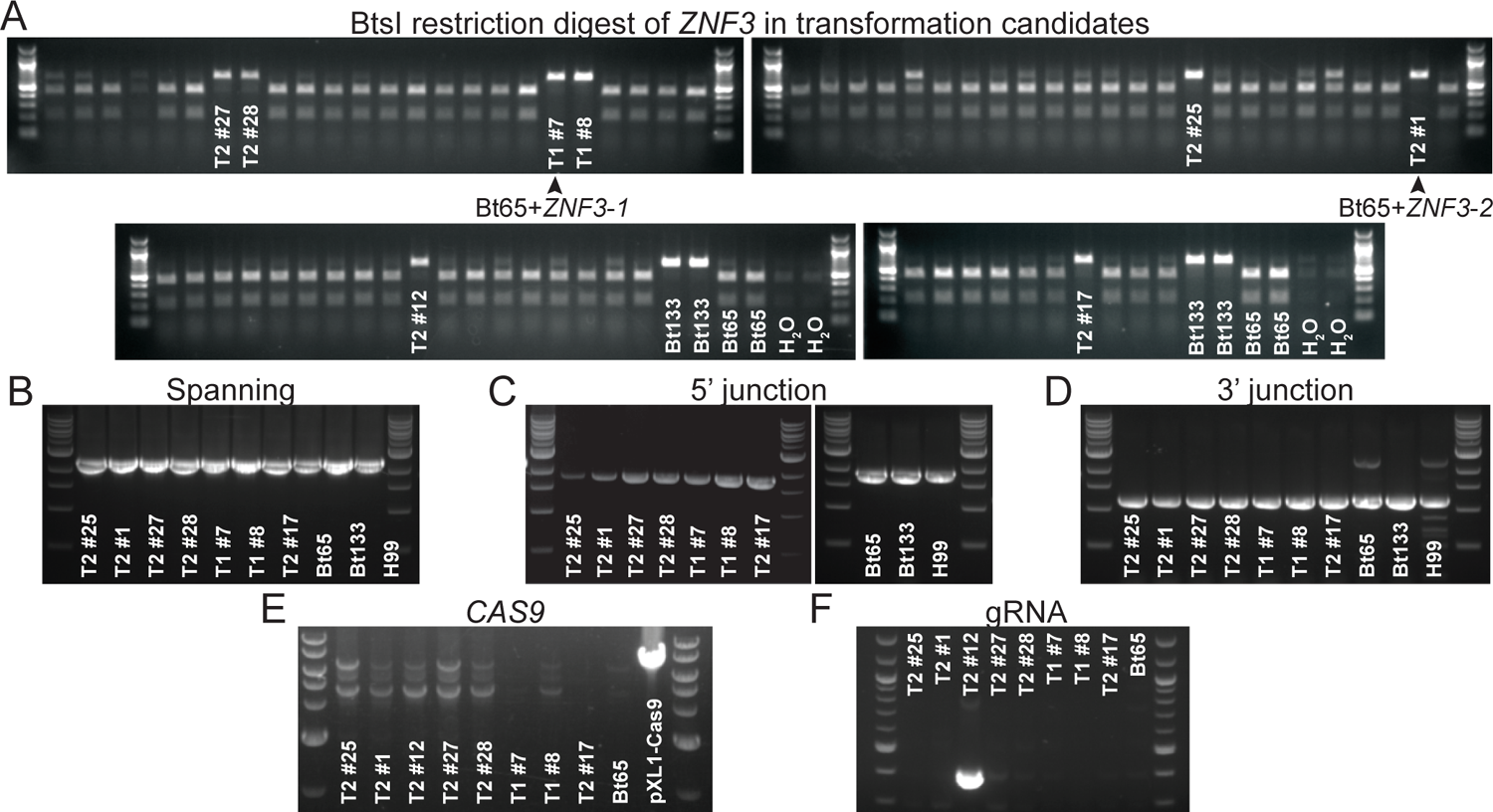
Identification and confirmation of correct Bt65+*ZNF3* transformants. **(A)** BtsI restriction enzyme digestion of *ZNF3* PCR products from nourseothricin-resistant transformants and controls (primers SJP186/187). **(B)** PCR amplification of the ZNF3 allele using primers outside of the Bt133 *ZNF3* allele used for homologous recombination (primers SJP208/209). PCR amplification to ensure correct integration of the **(C)** 5’ and **(D)** 3’ ends of the Bt133 *ZNF3* allele at the endogenous *ZNF3* locus (primers SJP208/187, and SJP186/209, respectively). PCR to ensure neither **(E)** *CAS9* nor **(F)** the gRNA constructs were integrated into the transformants (primers JOHE41657/45812 and JOHE50451/50452, respectively).

**Supplementary Figure S21.**
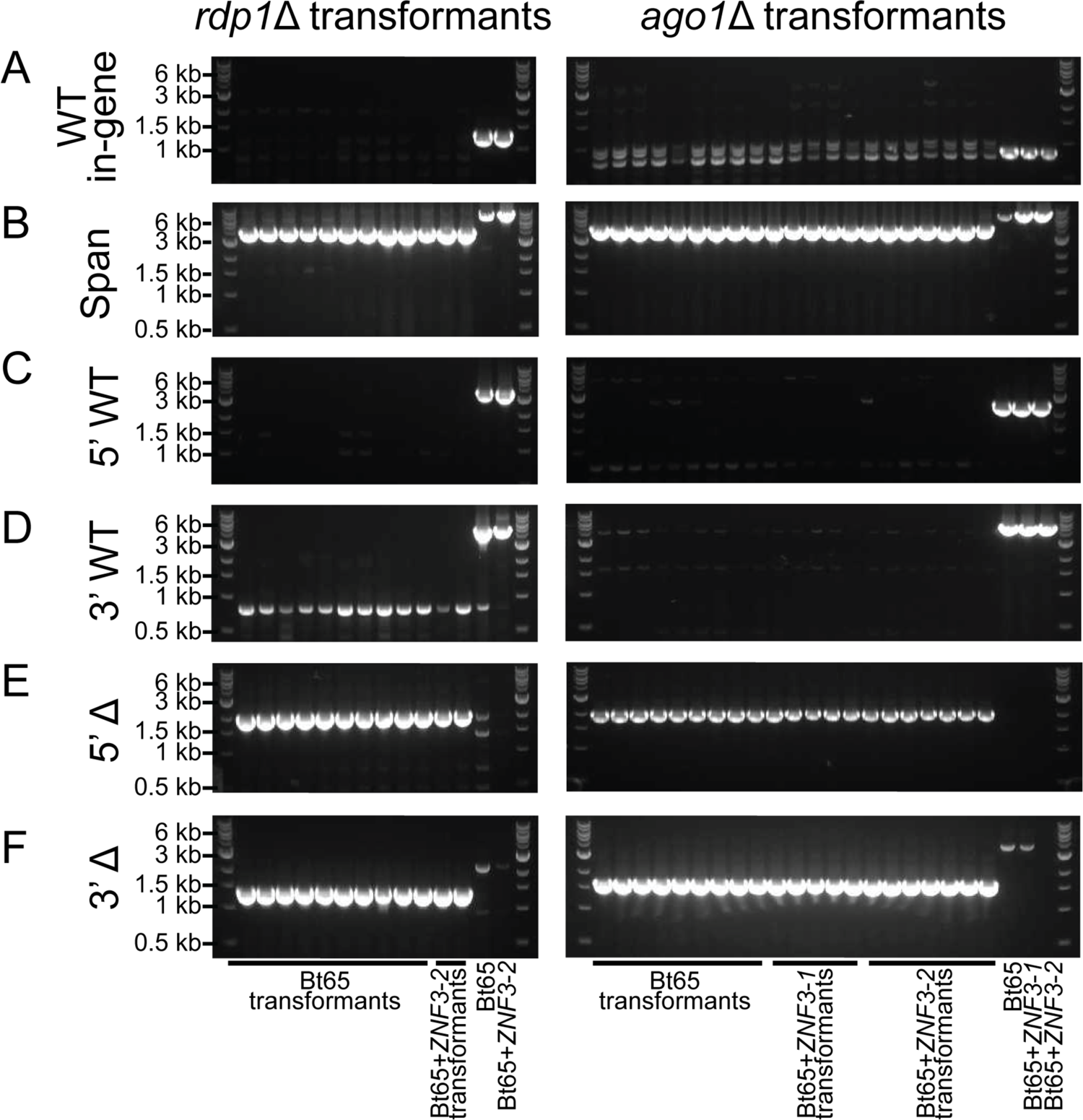
Confirmation of correct *rdp1*Δ and *ago1*Δ transformants. **(A)** PCR amplification of the *RDP1* and *AGO1* wild-type (WT) alleles using primers internal to the indicated ORF (primers SJP146/147 & SJP247/248, respectively). **(B)** Confirmation that only a single deletion construct was integrated at the *RDP1* (SJP144/145) and *AGO1* (SJP245/246) endogenous loci. PCR amplification to confirm absense of the **(C)** 5’ (*RDP1*: SJP144/147; *AGO1*: SJP245/248) and **(D)** 3’ ends (*RDP1*: SJP146/SJP145; *AGO1*: SJP247/246) of the indicated wild-type alleles. PCR amplification to confirm presence of the **(E)** 5’ (*rdp1*: SJP144/141; *ago1*: SJP245/242) and **(F)** 3’ ends (*rdp1*: SJP140/SJP145; *ago1*: SJP241/246) of the indicated deletion (Δ) constructs.

**Supplementary Figure S22.**
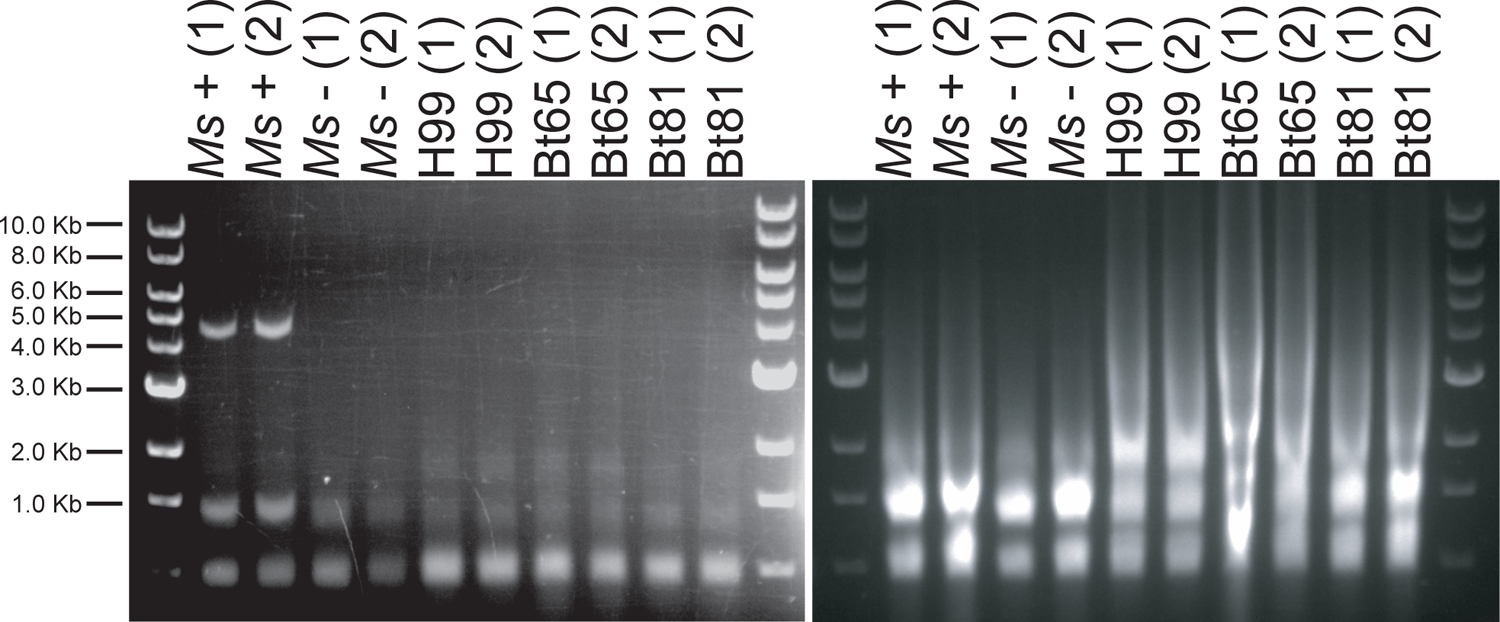
Enrichment for dsRNA does not identify any fragments likely to be dsRNA mycoviruses. Pictured on the left are RNA samples following LiCl enrichment for dsRNA run on a 1% agarose gel. Total RNA prior to dsRNA enrichment is pictured on the right on a 1% agarose gel. *Ms*+ is a *Malassezia sympodialis* strain that harbors a dsRNA virus, and *Ms*-is a congenic virus-cleared strain^69^. Two biological replicates for all samples are shown and labeled (1) and (2). The TriDye 1 kb DNA ladder (NEB) was used to estimate RNA fragment sizes.

## Supplementary Table Legends

**Supplementary Table S1.** Strains included in preliminary screen of SDC isolates for hypermutation phenotype.

**Supplementary Table S2.** Genetic variants and predicted changes in genes within QTL between H99 and Bt65.

**Supplementary Table S3.** (A) Centromere lengths in H99, Bt65, Bt81, Bt89, and Bt133, and (B) one-way ANOVA and Tukey’s HSD post hoc statistical tests for differences in mean centromere length.

**Supplementary Table S4.** sRNA analysis

**Supplementary Table S5.** Strains used in this study.

**Supplementary Table S6.** Mutation rates and 95% confidence intervals for all fluctuation assays.

**Supplementary Table S7.** Fluctuation assay data and calculated mutation frequencies

**Supplementary Table S8.** Oligonucleotides used in this study.

**Supplementary Table S9.** Cnl1 insertion sequences in PCR products.

**Supplementary Table S10.** Aneuploid, diploid, and clonal Bt65 x H99 F_1_ progeny.

