## Supplementary Table S3 for "Rampant transposition following RNAi loss causes hypermutation and antifungal drug resistance in clinical isolates of a human fungal pathogen"

**Table S3. (A) Centromere lengths in H99, Bt65, Bt81, Bt89, and Bt133, and (B) one-way ANOVA and Tukey’s HSD post hoc statistical test for differences in mean centromere length.**

**(A)**

|  | ***CEN* length (bp)** | | | | |
| --- | --- | --- | --- | --- | --- |
| **Centromere** | **H99** | **Bt65** | **Bt81** | **Bt89** | **Bt133** |
| *CEN1* | 37112 | 62504 | 62143 | 62573 | 62484 |
| *CEN2* | 60923 | 42388 | 42239 | 31179 | 34652 |
| *CEN3* | 49157 | 22835 | 22775 | 27206 | 26395 |
| *CEN4* | 43660 | 36285 | 36120 | 41252 | 51592 |
| *CEN5* | 42322 | 26280 | 26226 | 26255 | 26315 |
| *CEN6* | 41316 | 40179 | 39573 | 21109 | 34231 |
| *CEN7* | 54156 | 28455 | 28402 | 33812 | 34491 |
| *CEN8* | 49595 | 43749 | 43507 | 30907 | 30858 |
| *CEN9* | 37696 | 63297 | 62619 | 46791 | 65408 |
| *CEN10* | 44840 | 30765 | 30545 | 28835 | 35270 |
| *CEN11* | 64918 | 33348 | 32643 | 32893 | 52923 |
| *CEN12* | 31445 | 31772 | 31558 | 31776 | 31794 |
| *CEN13* | 52717 | 44526 | 44449 | 45915 | 50478 |
| *CEN14* | 52499 | 57218 | 56860 | 38514 | 39859 |
| Average length | 47311.14 | 40257.07 | 39975.64 | 35644.07 | 41196.43 |

**(B)**

**One-way ANOVA**

|  | **Degrees of Freedom** | **Sum of Squares** | **Mean Square** | **F value** | **p-Value** |
| --- | --- | --- | --- | --- | --- |
|  | 4 | 981127370 | 245281842 | 1.736 | 0.153 |
| **Residuals** | 65 | 9181816714 | 141258719 |  |  |

**Tukey’s HSD**

|  |  |  | **95% Confidence Intervals** | |  |
| --- | --- | --- | --- | --- | --- |
| **Strain** | **Strain** | **Differences** | **Lower** | **Upper** | **p-Value** |
| H99 | Bt65 | -7054.0714 | -19658.381 | 5550.2379 | 0.5215382 |
|  | Bt81 | -7335.5000 | -19939.809 | 5268.8093 | 0.4822454 |
|  | Bt89 | -11667.0714 | -24271.381 | 937.2379 | 0.0827166 |
|  | Bt133 | -6114.7143 | -18719.024 | 6489.5950 | 0.6543001 |
| Bt65 | Bt81 | -281.4286 | -12885.738 | 12322.8808 | 0.9999964 |
|  | Bt89 | -4613.0000 | -17217.309 | 7991.3093 | 0.8420273 |
|  | Bt133 | 939.3571 | -11664.952 | 13543.6665 | 0.9995630 |
| Bt81 | Bt89 | -4331.5714 | -16935.881 | 8272.7379 | 0.8700895 |
|  | Bt133 | 1220.7857 | -11383.524 | 13825.0950 | 0.9987719 |
| Bt89 | Bt133 | 5552.3571 | -7051.952 | 18156.6665 | 0.7304893 |
