## Supplementary Table S5 for "Rampant transposition following RNAi loss causes hypermutation and antifungal drug resistance in clinical isolates of a human fungal pathogen"

**Table S5. Strains used in this study.**

| **Strain** | **Strain number** | **Genotype** | **Comment** | **Source** |
| --- | --- | --- | --- | --- |
| H99α | JOHE4413 | *MAT*α | H99F isolate | [57] |
| *msh2*Δ | SJP241 | *MAT*α *msh2*Δ::*NAT* | From 2015 Madhani deletion collection; KN99α genetic background | [52] |
| PMHc1051 | SJP454 | VNBII *MAT*α |  | John Perfect |
| NRHc5014.ENR | SJP228 | VNBII *MAT*α |  | John Perfect |
| Bt133 | JOHE5131 | VNBII *MAT***a** | Also known as MMRL2989 | Tom Mitchell |
| Bt65 | JOHE3173 | VNBII *MAT***a** | Also known as MMRL2925 | Tom Mitchell |
| Bt81 | JOHE5082 | VNBII *MAT***a** | Also known as MMRL2940 | Wiley Schell |
| Bt89 | JOHE5090 | VNBII *MAT*α | Also known as MMRL2948 | Wiley Schell |
| Bt98 | JOHE5098 | VNBII *MAT*α | Also known as MMRL2956 | Wiley Schell |
| Bt84 | JOHE5085 | VNBII *MAT*α | Also known as MMRL2943 | Wiley Schell |
| Bt40 | JOHE5040 | VNBII *MAT*α | Also known as MMRL2898 | Wiley Schell |
| PMHc1035 | SJP234 | VNBII *MAT*α |  | John Perfect |
| Bt146 | JOHE5142 | VNBII *MAT*α | Also known as MMRL3000 | Wiley Schell |
| Bt115 | JOHE5114 | VNBII *MAT*α | Also known as MMRL2972 | Wiley Schell |
| Bt76 | JOHE5077 | VNBII *MAT*α | Also known as MMRL2935 | Wiley Schell |
| *crg1*Δ | JOHE4514 | *MAT*α *ura5^-^ crg1*::*URA5* | F99 genetic background | [56] |
| Bt65+*ZNF3*-*1* | SJP581 | *MAT***a** *ZNF3* | Bt65 genetic background, *ZNF3* allele from Bt133, independent from SJP584 | This study |
| Bt65+*ZNF3*-*2* | SJP584 | *MAT***a** *ZNF3* | Bt65 genetic background, *ZNF3* allele from Bt133, independent from SJP581 | This study |
| *rdp1*Δ | JOHE6840 | *MAT*α *rdp1*::*NEO* | H99 genetic background | [35] |
| *znf3*Δ | JOHE12984 | *MAT*α *znf3*::*NAT* | H99 genetic background; MF195 | [38] |
| *ago1*Δ | JOHE4933 | *MAT*α *ago1*::*NAT* | H99 genetic background | [14] |
| Bt65 *rdp1*Δ | SJP667 | *MAT***a** *rdp1*::*NAT* | Bt65 genetic background | This study |
| Bt65 *ago1*Δ | SJP672 | *MAT***a** *ago1*::*NAT* | Bt65 genetic background | This study |
| Bt65+*ZNF3*-*1 ago1*Δ | SJP676 | *MAT***a** *ZNF3 ago1*::*NAT* | Bt65+*ZNF3*-*1* genetic background | This study |
| Bt65+*ZNF3*-*2 rdp1*Δ | SJP671 | *MAT***a** *ZNF3 rdp1*::*NAT* | Bt65+*ZNF3*-*2* genetic background | This study |
| Bt65+*ZNF3*-*2 ago1*Δ | SJP678 | *MAT***a** *ZNF3 ago1*::*NAT* | Bt65+*ZNF3*-*2* genetic background | This study |
| *Ms* + | JOHE10102 | MsMV1 infected | Strain KS012; *Malassezia sympodialis* clinical isolate with dsRNA virus | [69] |
| *Ms* - | JOHE18529 | MsMV1 temperature-cured | Congenic strain of KS012; *Malassezia sympodialis* clinical isolate cleared of dsRNA virus by 37°C passage | [69] |
| pSDMA57 | Plasmid #323 | Amp^R^ *NEO* | *E. coli*; *C. neoformans* safe haven plasmid SH1-NEO | [94] |
| pXL1-Cas9-HYG | Plasmid #488 | Kan^R^ G418^R^ | *E. coli*; TRACE plasmid | [58] |
| pYF511 | Plasmid #489 | Amp^R^ | Upgraded all-in-one Cas9 + sgRNA plasmid | [96] |
| pAI3 | Plasmid #263 | Kan^R^ NAT^R^ | *E. coli*; Actin promoter from JEC21, Nourseothricin acetyltransferase (synthetic, optimized for *Cryptococcus*), terminator *TRP1* from JEC21. | [95] |
| Bt65 F_1_  progeny #2 | SJP243 | *znf3* | H99α *crg1*Δ x Bt65**a**  F_1_ progeny; hypermutator | This study |
| Bt65 F_1_  progeny #3 | SJP244 | *znf3* | H99α *crg1*Δ x Bt65**a**  F_1_ progeny; hypermutator | This study |
| Bt65 F_1_  progeny #8 | SJP249 | *znf3* | H99α *crg1*Δ x Bt65**a**  F_1_ progeny; hypermutator | This study |
| Bt65 F_1_  progeny #11 | SJP382 | *ZNF3* | H99α *crg1*Δ x Bt65**a**  F_1_ progeny;  non-hypermutator | This study |
| Bt65 F_1_  progeny #12 | SJP253 | *ZNF3* | H99α *crg1*Δ x Bt65**a**  F_1_ progeny;  non-hypermutator | This study |
| Bt65 F_1_  progeny #13 | SJP383 | *ZNF3* | H99α *crg1*Δ x Bt65**a**  F_1_ progeny;  non-hypermutator | This study |
| Bt65 F_1_  progeny #14 | SJP384 | *ZNF3* | H99α *crg1*Δ x Bt65**a**  F_1_ progeny; non-hypermutator | This study |
| Bt65 F_1_  progeny #17 | SJP258 | *ZNF3* | H99α *crg1*Δ x Bt65**a**  F_1_ progeny; non-hypermutator | This study |
| Bt65 F_1_  progeny #18 | SJP259 | *ZNF3* | H99α *crg1*Δ x Bt65**a**  F_1_ progeny; non-hypermutator | This study |
| Bt65 F_1_  progeny #19 | SJP260 | *znf3* | H99α *crg1*Δ x Bt65**a**  F_1_ progeny; hypermutator | This study |
| Bt65 F_1_  progeny #20 | SJP261 | *ZNF3* | H99α *crg1*Δ x Bt65**a**  F_1_ progeny;  non-hypermutator | This study |
| Bt65 F_1_  progeny #21 | SJP262 | *ZNF3* | H99α *crg1*Δ x Bt65**a**  F_1_ progeny; non-hypermutator | This study |
| Bt65 F_1_  progeny #22 | SJP270 | *ZNF3* | H99α *crg1*Δ x Bt65**a**  F_1_ progeny; non-hypermutator | This study |
| Bt65 F_1_  progeny #23 | SJP271 | *znf3* | H99α *crg1*Δ x Bt65**a**  F_1_ progeny; hypermutator | This study |
| Bt65 F_1_  progeny #24 | SJP272 | *znf3* | H99α *crg1*Δ x Bt65**a**  F_1_ progeny; hypermutator | This study |
| Bt65 F_1_  progeny #25 | SJP273 | *ZNF3* | H99α *crg1*Δ x Bt65**a**  F_1_ progeny; non-hypermutator | This study |
| Bt65 F_1_  progeny #26 | SJP274 | *ZNF3* | H99α *crg1*Δ x Bt65**a**  F_1_ progeny; non-hypermutator | This study |
| Bt65 F_1_  progeny #27 | SJP275 | *znf3* | H99α *crg1*Δ x Bt65**a**  F_1_ progeny; hypermutator | This study |
| Bt65 F_1_  progeny #29 | SJP277 | *znf3* | H99α *crg1*Δ x Bt65**a**  F_1_ progeny; hypermutator | This study |
| Bt65 F_1_  progeny #30 | SJP278 | *ZNF3* | H99α *crg1*Δ x Bt65**a**  F_1_ progeny;  non-hypermutator | This study |
| Bt65 F_1_  progeny #32 | SJP280 | *znf3* | H99α *crg1*Δ x Bt65**a**  F_1_ progeny; hypermutator | This study |
| Bt65 F_1_  progeny #33 | SJP281 | *ZNF3* | H99α *crg1*Δ x Bt65**a**  F_1_ progeny;  non-hypermutator | This study |
| Bt65 F_1_  progeny #34 | SJP282 | *znf3* | H99α *crg1*Δ x Bt65**a**  F_1_ progeny; hypermutator | This study |
| Bt65 F_1_  progeny #36 | SJP284 | *ZNF3* | H99α *crg1*Δ x Bt65**a**  F_1_ progeny; non-hypermutator | This study |
| Bt65 F_1_  progeny #37 | SJP285 | *ZNF3* | H99α *crg1*Δ x Bt65**a**  F_1_ progeny; non-hypermutator | This study |
| Bt65 F_1_  progeny #39 | SJP287 | *znf3* | H99α *crg1*Δ x Bt65**a**  F_1_ progeny;  hypermutator | This study |
| Bt65 F_1_  progeny #41 | SJP289 | *znf3* | H99α *crg1*Δ x Bt65**a**  F_1_ progeny; hypermutator | This study |
| Bt65 F_1_  progeny #44 | SJP292 | *znf3* | H99α *crg1*Δ x Bt65**a**  F_1_ progeny; hypermutator | This study |
| Bt81 F_1_  progeny #1 | SJP296 | *znf3* | H99α *crg1*Δ x Bt81**a**  F_1_ progeny | This study |
| Bt81 F_1_  progeny #2 | SJP297 | *ZNF3* | H99α *crg1*Δ x Bt81**a**  F_1_ progeny | This study |
| Bt81 F_1_  progeny #3 | SJP298 | *ZNF3* | H99α *crg1*Δ x Bt81**a**  F_1_ progeny | This study |
| Bt81 F_1_  progeny #4 | SJP299 | *ZNF3* | H99α *crg1*Δ x Bt81**a**  F_1_ progeny | This study |
| Bt81 F_1_  progeny #5 | SJP300 | *ZNF3* | H99α *crg1*Δ x Bt81**a**  F_1_ progeny | This study |
| Bt81 F_1_  progeny #6 | SJP301 | *ZNF3* | H99α *crg1*Δ x Bt81**a**  F_1_ progeny | This study |
| Bt81 F_1_  progeny #7 | SJP302 | *ZNF3* | H99α *crg1*Δ x Bt81**a**  F_1_ progeny | This study |
| Bt81 F_1_  progeny #8 | SJP303 | *ZNF3* | H99α *crg1*Δ x Bt81**a**  F_1_ progeny | This study |
| Bt81 F_1_  progeny #9 | SJP304 | *ZNF3* | H99α *crg1*Δ x Bt81**a**  F_1_ progeny | This study |
| Bt81 F_1_  progeny #10 | SJP305 | *znf3* | H99α *crg1*Δ x Bt81**a**  F_1_ progeny | This study |
| Bt81 F_1_  progeny #11 | SJP306 | *ZNF3* | H99α *crg1*Δ x Bt81**a**  F_1_ progeny | This study |
| Bt81 F_1_  progeny #12 | SJP307 | *ZNF3* | H99α *crg1*Δ x Bt81**a**  F_1_ progeny | This study |
| Bt81 F_1_  progeny #13 | SJP308 | *znf3* | H99α *crg1*Δ x Bt81**a**  F_1_ progeny | This study |
| Bt81 F_1_  progeny #14 | SJP309 | *ZNF3* | H99α *crg1*Δ x Bt81**a**  F_1_ progeny | This study |
| Bt81 F_1_  progeny #15 | SJP310 | *ZNF3* | H99α *crg1*Δ x Bt81**a**  F_1_ progeny | This study |
| Bt81 F_1_  progeny #17 | SJP312 | *ZNF3* | H99α *crg1*Δ x Bt81**a**  F_1_ progeny | This study |
| Bt81 F_1_  progeny #18 | SJP313 | *ZNF3* | H99α *crg1*Δ x Bt81**a**  F_1_ progeny | This study |
| Bt81 F_1_  progeny #34 | SJP329 | *znf3* | H99α *crg1*Δ x Bt81**a**  F_1_ progeny | This study |
