## Supplementary Table S8 for "Rampant transposition following RNAi loss causes hypermutation and antifungal drug resistance in clinical isolates of a human fungal pathogen"

**Table S8. Oligonucleotides used in this study.**

| **Primer** | **Sequence (5’ to 3’)** | **Comment** |
| --- | --- | --- |
| JOHE42462/ZC7 | ACAGTCACGAGCCCTGAAAC | *C. neoformans* (H99) *FRR1* F |
| JOHE42463/ZC8 | AGTCGGAGGTTTGGACAGTG | *C. neoformans* (H99) *FRR1* R |
| JOHE43186/ZC19 | CGATAACATCTGCGACGAAA | *C. neoformans* (H99) *URA5* F |
| JOHE43187/ZC20 | TAGCCTCCTTTGTCGCTTCA | *C. neoformans* (H99) *URA5* R |
| JOHE44472/SJP43 | CCAGCTCGTACCTTGCTCTG | *C. neoformans* (H99) *URA3* F |
| JOHE44473/SJP44 | CAGGGCTGTTGTCTCGTAGC | *C. neoformans* (H99) *URA3* R |
| JOHE45507/SJP107 | ACGTGACTGGATTGCTGATTG | *F primer upstream of FUR1 5'UTR* |
| JOHE45508/SJP108 | GACTCAACCTTGCTTTCCCATC | *R primer downstream of FUR1 3'UTR* |
| JOHE45509/SJP109 | GAGGAAGGTGAGTTTTCGAATG | *F primer FUR1 in gene* |
| JOHE45510/SJP110 | AATGTCATCGGGCAACTTAGC | *R primer FUR1 in gene* |
| JOHE45511/SJP111 | ACGCAGAGAGAAAAGCTCCAG | F primer upstream of *UXS1* 5’UTR |
| JOHE45512/SJP112 | CGCAAAATCAACTCGTCATTTC | R primer downstream of *UXS1* 3’ UTR |
| JOHE45513/SJP113 | ACCGTCCTCGACAACTTCTTC | F primer *UXS1* in gene |
| JOHE45514/SJP114 | AGTCGTGGACGTATTGGAAGG | R primer *UXS1* in gene |
| JOHE50451 | GCTCATGGATCCTTTGCATTAGAACTAAAAACAAAGCA | gRNA construct: U6 promoter F |
| JOHE50452 | GATCATCCGCGGtaaaacaaaaaagcaccgactcggtgcc | gRNA construct: gRNA scaffold R |
| JOHE50954/SJP200 | CGGCTAGTGAAGAACGAACTTG | Bt133 *ZNF3* allele for CRISPR-mediated gene replacement F |
| JOHE50955/SJP201 | GCATCAAGAAAGCAGCATTTG | Bt133 *ZNF3* allele for CRISPR-mediated gene replacement R |
| JOHE50952/SJP198 | caaagtggaaattgcacatacaccggcagggtatactgttgATCGAGCAGCAACAGCAGTGgttttagagctagaaatagc | gRNA construct: Bt65 *ZNF3* gRNA with U6 and scaffold homology F |
| JOHE50953/SJP199 | gctatttctagctctaaaacCACTGCTGTTGCTGCTCGATcaacagtataccctgccggtgtatgtgcaatttccactttg | Bt65 *ZNF3* gRNA with U6 and scaffold homology F |
| JOHE46609 | AGTGCTGTGGTGAAAGAGAT GTTTTAGAGCTAGAAATAGCAAG | Safe Haven 1 gRNA with homology to sgRNA scaffold F |
| JOHE46608 | ATCTCTTTCACCACAGCACTCAACAGTATACCCTGCCGGTG | Safe Haven 1 gRNA with homology to U6 R |
| JOHE45941 | ggtgacgctgtgagagtgg | *CAS9* F |
| JOHE45942 | gggcccctcttcacgtgg | *CAS9* R |
| JOHE41657 | CATGCATCTAGGTCTAGAAACC | *GPD1* promoter F |
| JOHE45812 | ACTGGCCGTCGTTTTACCTCTTCACGTGGACGCTCC | *GPD1* promoter end + *NAT* R |
| JOHE50145 | TTGGCTACCCGTGATATTGC | *NEO* confirmation F |
| JOHE50177 | TTGGATCCTCAATTGTCTCCT | Safe Haven 1 integration upstream confirmation F |
| JOHE50006/SJP186 | GAACAGGAGGCACGAAGC | *ZNF3* upstream 5' UTR F |
| JOHE50007/SJP187 | CCTCTCAGTAGGTGCGGAAG | *ZNF3* internal R |
| JOHE50294/SJP189 | GACTCACATGCAGCCAGCAG | *ZNF3* upstream 5' UTR (outside Bt133 complementation allele) F |
| JOHE50295/SJP190 | GCCTCTCGCTGAGCTTGAAC | *ZNF3* downstream 3’ UTR (outside Bt133 complementation allele) R |
| JOHE23951/XW145 | CAAGAACCTCAAGGTGGTAGAATTA | H99 FKBP12 5' F |
| JOHE23952/XW146 | AGGAATTTCAATCAGCTAAATACCC | H99 FKBP12 5' R |
| JOHE45581/SJP127 | CTATGAAATATGCGCACCTTCG | *FRR1* (CNAG_03682) in gene R |
| JOHE45582/SJP128 | CGAAGGTGCGCATATTTCATAG | *FRR1* (CNAG_03682) in gene F |
| SJP135 | CATCCGCTTTGTGAGTGTGTG | Bt65 insertion R-internal primer |
| SJP136 | AGCGCAGGAAGTAGAACCAG | Bt65 insertion-internal F primer |
| SJP137 | CATAGCAAAAGGTCGCCATC | *FRR1* in-gene R primer for insertion sequencing |
| SJP148 | CTCAGTCGTCAAGTTCCTCACC | Bt65 insertion-internal F primer #2 |
| SJP149 | CTTGTCGAGCCAGGAGAGAC | Bt65 insertion-internal R primer #2 |
| JOHE51176/SJP208 | TGTTGTCGCATCCACAGCTAC | *ZNF3* (CNAG_02700) F outside Bt133 allele used for CRISPR-mediated replacement |
| JOHE51177/SJP209 | TTGCAGACGGGAAGCTCAATC | *ZNF3* (CNAG_02700) R outside Bt133 allele used for CRISPR-mediated replacement |
| SJP138 | CTCAATGTTGAGGACCTTGACC | *RDP1* (CNAG_03466) 5' fragment F, starts 966bp upstream ATG |
| SJP139 | CTGGCCGTCGTTTTACTTGACGGGAATTTCAGC | *RDP1* (CNAG_03466) 5' fragment R, starts immediately upstream ATG, split with *NAT* |
| SJP140 | GCTGAAATTCCCGTCAAGTAAAACGACGGCCAG | *NAT* F primer to delete *RDP1* (CNAG_03466), split with CNAG_03466 |
| SJP141 | CGAAGAAGAAGTCGACAGCAGGAAACAGCTATGAC | *NAT* R primer to delete *RDP1* (cnag_03466), split with *RDP1* |
| SJP142 | GTCATAGCTGTTTCCTGCTGTCGACTTCTTCTTCG | *RDP1* (CNAG_03466) 3' fragment F, starts 223bp downstream of STOP, split with *NAT* |
| SJP143 | GAATGTGAATCCAGCTTCTTG | *RDP1* (CNAG_03466) 3' fragment R, starts 1030bp downstream of STOP |
| SJP144 | CATTTCGCATGACGAGTTTTTG | *RDP1* (CNAG_03466) 5' F outside of deletion construct |
| SJP145 | AGTTGGCTGTGGTTGTCATTC | *RDP1* (CNAG_03466) 3' R outside of deletion construct |
| SJP146 | GCTCCAGGTGATACAGCAGAG | *RDP1* (CNAG_03466) internal F |
| SJP147 | TCCGTTACTGCGTTATTCCAC | *RDP1* (CNAG_03466) internal R |
| JOHE52135/SJP250 | accggcagggtatactgttgAGATGGGGTTGGATGTATGGgttttagagctagaaatagc | Bt65 *RDP1* (CNAG_03466) gRNA #2 with U6 and scaffold homology F |
| JOHE52136/SJP251 | accggcagggtatactgttgAAACGCAGAATACTCACAGGgttttagagctagaaatagc | Bt65 *RDP1* (CNAG_03466) gRNA #3 with U6 and scaffold homology F |
| JOHE52137/SJP252 | accggcagggtatactgttgGTAGTTACCGAACCTAGCGGgttttagagctagaaatagc | Bt65 *AGO1* (CNAG_04609) gRNA #1 with U6 and scaffold homology F |
| JOHE52138/SJP253 | accggcagggtatactgttgGTTCTCCAAGCTTTCCTTTGgttttagagctagaaatagc | Bt65 *AGO1* (CNAG_04609) gRNA #2 with U6 and scaffold homology F |
| JOHE52103/SJP239 | CATTGAGCTGAATCCGAACAG | *AGO1* (CNAG_04609) 5' fragment F, starts 1060bp upstream ATG |
| JOHE52104/SJP240 | CTGGCCGTCGTTTTAAGGAAGGAAGAAATTGG | *AGO1* (CNAG_04609) 5' fragment R, starts 111bp upstream ATG, split with *NAT* |
| JOHE52105/SJP241 | CCAATTTCTTCCTTCCTTAAAACGACGGCCAG | *NAT* F primer to delete *AGO1* (CNAG_04609), split with *AGO1* |
| JOHE52106/SJP242 | CTTTCGCATTTCGATCCAGGAAACAGCTATGAC | *NAT* R primer to delete *AGO1* (CNAG_04609), split with *AGO1* |
| JOHE52107/SJP243 | GTCATAGCTGTTTCCTGGATCGAAATGCGAAAG | *AGO1* (CNAG_04609) 3' fragment F, starts 69bp downstream of STOP, split with *NAT* |
| JOHE52108/SJP244 | TCATCCGCAACTTTAGCTTTG | *AGO1* (CNAG_04609) 3' fragment R, starts 1,096bp downstream of STOP |
| JOHE52109/SJP245 | CACCCCTGCCTTACCTTTAAC | *AGO1* (CNAG_04609) 5' junction check F |
| JOHE52110/SJP246 | ATTAGCCGAAGCTGAACTTGC | *AGO1* (CNAG_04609) 3' junction check R |
| JOHE52111/SJP247 | ATTGAGATCAACCCCGTCATC | *AGO1* (CNAG_04609) in-gene F |
| JOHE52112/SJP248 | GAAGGTGCGAGACCAAGAAC | *AGO1* (CNAG_04609) in-gene R |
