## Supplementary Table S10 for "Rampant transposition following RNAi loss causes hypermutation and antifungal drug resistance in clinical isolates of a human fungal pathogen"

**Table S10. Aneuploid, diploid, and clonal Bt65 x H99 F_1_ progeny.**

| Strain | Description | Retained for QTL analysis |
| --- | --- | --- |
| S13 | Chromosome 11, duplication | yes |
| S14 | Chromosome 11 and Chromosome 13, heterozygotic aneuploidy | no |
| S20 | Chromosome 13, heterozygotic aneuploidy | yes |
| S25 | Heterozygotic diploid | no |
| S34 | Chromosome 4, partial duplication | yes |
| S41 | Chromosome 13, partial duplication | no |
| S12 | Clone of strain, S11 | no |
| S41 | Clone of strain, S23 | no |
